# Dynamic Exchange of Bacteria and Carbapenem Resistance Genes between Sewer Biofilms and Wastewater

**DOI:** 10.64898/2026.05.20.726639

**Authors:** Ean Warren, William J. Brazelton, Sydney Fusco, James VanDerslice, L. Scott Benson, Windy Tanner, Jennifer Weidhaas

## Abstract

Sewer biofilms represent dynamic interfaces for exchange of bacteria and antibiotic resistance genes between biofilms and the overlying wastewater. Using inline, biofilm reactors, the movement of bacteria and 16S rRNA and carbapenemase genes (*bla*_KPC_, *bla*_VIM_, *bla*_NDM_, *bla*_OXA-48-like_, and *bla*_IMP_) between wastewater and sewer biofilms was investigated. Established, complex biofilms without these β-lactamase (*bla*) genes, absorbed resistant bacteria within two minutes of exposure to high concentrations of resistant cultures in lab settings. Carbapenem-resistant organisms from these high-concentration source biofilms transferred to downstream biofilms over 60 minutes of representative sewer shear flows. Mass balances of bacteria and genes in biofilms versus wastewater under representative shear flow showed that biofilms exposed to resistant cultures contributed more to the wastewater than to the downstream biofilms. In field studies, established, complex biofilms without target carbapenem-resistant bacteria and genes from wastewater within hours and then stabilized between 2 to 15 days, not varying by more than 0.5 MPN/cm^2^ or 0.5 log gene copies (GC)/cm^2^. In contrast, metagenomic profiles of the bacterial community species continued to change up to 21 days. Established biofilms with resistant bacteria and genes exposed to tertiary-treated wastewater without target carbapenemase genes or meropenem antibiotics did not lose resistant genes or bacteria over nine days of exposure (i.e., < 1 log GC/cm^2^ reduction). Results show that sewer biofilms contribute to the resistance-gene signal found in sewer wastewater by absorbing and releasing bacteria and genes. Consideration of sewer biofilm dynamics is essential for more accurately interpreting wastewater bacterial concentrations in wastewater-based epidemiology studies.

**GRAPHICAL ABSTRACT:** 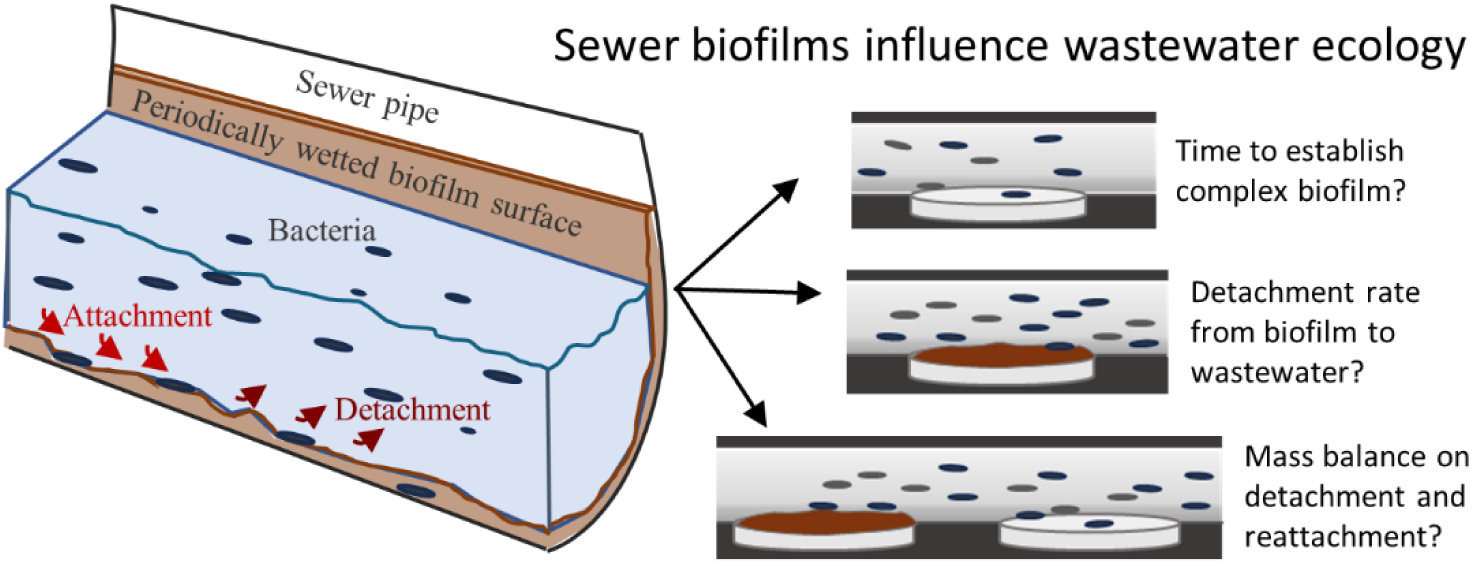

## 1.0 INTRODUCTION

Biofilms form under most natural and engineered conditions, including in the human body, on environmental surfaces, around sink drains, and in sewer collection pipes. Biofilms are complex structures of many microorganisms and extracellular polymeric substances (EPS), nucleic acids, polysaccharides, lipids, and other organic and inorganic compounds [1–3]. The biofilm structure provides microorganisms with protection against toxins, including antibiotics, protozoan grazers, and external physical and chemical stressors [4]. Biofilms in sewers and the overlying wastewater constantly exchange bacteria and genes and are influenced by various factors such as shear stress, total suspended solids, antimicrobial concentrations, and predation. The influence of sewer biofilms on wastewater-based epidemiology (WBE) (i.e., evaluating community health by analyzing wastewater) has largely been overlooked and current models assume that wastewater signals reflect only real-time human shedding of biomarkers [5–8]. From a clinical standpoint, failure to account for biofilm-mediated storage and release could lead to erroneous attribution of persistent resistance signals to ongoing patient transmission, potentially triggering unnecessary infection control responses or obscuring true outbreak timing [5, 7].

Sewer collection systems are a complex hydraulic, chemical, and biological network of wastewater from municipal and industrial sources with varying chemical and biological constituents [9]. Sewer systems from a single building are typically only partially filled with wastewater and have varying flow rates from fast, turbulent flows after toilet flushing to a trickle depending on the time of day or the number of sources contributing to the sewer. Sewer biofilms develop on the sides of the collection pipes and are influenced by the time-varying aerobic or anoxic conditions [10] and the variable flow rates. The sewer biofilm structure protects the biofilm community against bacterial toxins, including antibiotics, protozoan grazers, and external shocks such as temperature or chemical changes. Shear stress from the flowing liquid influences the types of cells that can accumulate and biofilm structural integrity [11–13]. The sewer biofilm structure acts as a miniature community with communication, protection, consumption, waste removal, replication, and gene transfer, including antimicrobial resistance gene (ARG) acquisition, occurring among the members.

Antimicrobial resistance (AMR) is a growing public health threat, responsible for more than 35,000 deaths each year in the United States and over 1.2 million worldwide [14, 15]. By 2050, deaths linked to AR are projected to rise sharply, positioning it as one of the leading causes of mortality worldwide [16]. One such emerging AMR threat is carbapenemase-producing organisms (CPOs), that actively express carbapenemase enzymes and carry functional carbapenemase genes. CPOs can shed a variety of carbapenemase genes into wastewater including *bla*_IMP_, *bla*_KPC_, *bla*_NDM_, *bla*_OXA-48-like_, and *bla*_VIM_. Further, carbapenem-resistant organisms (CROs), which encompass a broader group of organisms that may resist carbapenem antibiotics through non-enzyme mechanisms such as efflux pumps [17] and not carry carbapenemase genes at all, can also be present in wastewater. Investigating both CROs and carbapenemase genes in wastewater and sewer biofilms allows for interpretation of whether resistance arises from carbapenemase enzyme production or other mechanisms. These findings are clinically relevant because carbapenem-resistant Enterobacterales are strongly associated with healthcare settings, where colonized or infected patients may shed organisms prior to clinical detection. Wastewater surveillance has therefore been proposed as an adjunct to traditional infection control surveillance, particularly for early warning of emerging resistance in hospitals [18, 19].

Wastewater-based surveillance can be used to monitor both carbapenemase genes and CROs in sewers, such as from hospitals. Long-term monitoring of hospital sewers in Europe [20] and the United States [21–23] has shown that carbapenemase genes are consistently detected. Because carbapenemase genes can persist and vary over time in hospital wastewater, distinguishing true outbreaks of CPO infections from background noise is challenging. This problem is compounded by biofilms in sink drains, p-traps, and sewers, which may contribute carbapenemase genes or CRO to overflowing wastewater. These factors highlight the importance of considering CROs and carbapenemase genes in biofilms when utilizing WBE [7]. While some studies have focused on carbapenemase genes and CROs in wastewater and sewers [24–26], the movement of these organisms or genes in and out of sewer biofilms and their potential to confound WBE remain poorly understood. When colonized or infected individuals urinate and defecate into toilets, the added CRO, CPO, or carbapenemase genes in the sewer wastewater may partition to sewer biofilms, which may then act as long-term reservoirs for resistant bacteria because the biofilm structure protects cells from higher antibiotic concentrations and promotes gene exchange. The bacteria can also detach and colonize surfaces downstream or be measured in composite wastewater samples.

To understand the dynamics of CROs and carbapenemase genes in complex sewer biofilms and their impact on wastewater collected for WBE, we conducted five replicated experiments in laboratory and field settings, observing total and carbapenem-resistant bacteria as well as *bla*_KPC_, *bla*_VIM_, *bla*_NDM_, *bla*_OXA-48-like_, and *bla*_IMP_ genes. The studies were designed to simulate CROs and carbapenemase genes entering a building-scale sewer system from waste flushed from a colonized or infected individual. First, we determined how quickly a complex, stable biofilm could be established on sterile polyvinyl chloride (PVC) sewer pipe. Second, we evaluated how quickly CROs and carbapenemase genes attached to and detached from mature and complex biofilms. Third, we measured the loss of CRO and carbapenemase genes from mature and complex biofilms when exposed to water with low or no concentrations of CROs and carbapenemase genes and low to no antibiotics (i.e., removal of selective pressure). Finally, we measured the detachment and transport of CROs and carbapenemase genes from a biofilm source into the exposed wastewater and incorporation into downstream biofilms. The results of this study will help inform wastewater surveillance by determining the effects of sewer biofilms containing CROs and carbapenemase genes on downstream sewer wastewater.

## 2.0 MATERIALS AND METHODS

### 2.1 Materials

All reagents used in this study were of molecular biology grade unless otherwise specified. See Supplemental information for chemicals and sources.

### 2.2 Analytical methods

Since the treatment facility’s grit basin and effluent waters were used as reactor influent, meropenem concentrations were measured in each using filtration, solid phase extraction, and liquid chromatography–mass spectrometry (supplemental Table S1). Biofilm and grit chamber total and volatile solids were measured using gravimetric analysis. See supplemental information for method details.

### 2.3 Quantification of total and resistant bacteria

Biofilm bacteria on PVC coupons were measured by rinsing coupons with 10 mL of sterile 1X phosphorus-buffered saline (PBS) to remove loosely attached bacteria, then scraping across the diameter of the coupon twice with sterile pipette tips. Total bacterial counts were determined using Luria-Bertani (LB) in agar plates or most probable number assay (MPNs). Carbapenem-resistant bacteria were determined by MPN in LB augmented with carbapenem antibiotics (i.e., 8 mg/L meropenem or imipenem) or were determined with CHROMagar KPC plates. **Supplemental Table S2** contains a list of pure bacterial strains used in this study. More details are found in the supplemental information.

### 2.4 Quantification of carbapenemase genes, 16S rRNA, and metagenomes

Nucleic acid extraction from coupons, influent grit basin water, and effluent reactor water was performed by following a previously published method [27] with the exceptions found in supplemental information. Carbapenemase genes (i.e., *bla*_IMP_, *bla*_KPC_, *bla*_OXA-48-like_, *bla*_NDM_, *bla*_VIM_) and 16S rRNA were measured by quantitative polymerase chain reaction (qPCR) as has been described previously [22]. More details are presented in **supplemental Tables S3** to **S5**. The limits of detection for carbapenemase genes and bacteria on the coupons and in wastewater were 146 ± 92 gene copies (GC)/mL and 38 ± 41 colony-forming units (CFU)/mL for *bla*_VIM_, 284 ± 154 GC/mL and 25 ± 24 CFU/mL for *bla*_NDM_, 5486 ± 3137 GC/mL and 337 ± 341 CFU/mL for *bla*_IMP_ as described previously [22]. Metagenomic sequencing was used to characterize microbial communities and antimicrobial resistance genes. See supplemental information for method information.

### 2.5 Time to establish a stable biofilm from wastewater

Experiment 1 was designed to observe the development of a stable biofilm on PVC using wastewater as the reactor influent under realistic shear stress (**Table 1** and **Figure 1A**). A single, sterile inline reactor was deployed inside a temperature-controlled building at a wastewater treatment facility (WWTF). PVC coupons were initially sterilized by autoclave while reactors and tubing were sterilized with a 10% bleach solution and then rinsed with sterile deionized water. The reactor influent was grit basin wastewater (water quality shown in **supplemental Table S6**) continuously pumped from the surface of the basin. The reactors were inverted with the plugs on the bottom, ensuring a constant flow of water over the coupons and no air bubbles. During the three-week study period, the coupons were sampled in duplicate starting from the influent side and replaced with sterile coupons at the times indicated in **Table 1**.

**Figure 1.**
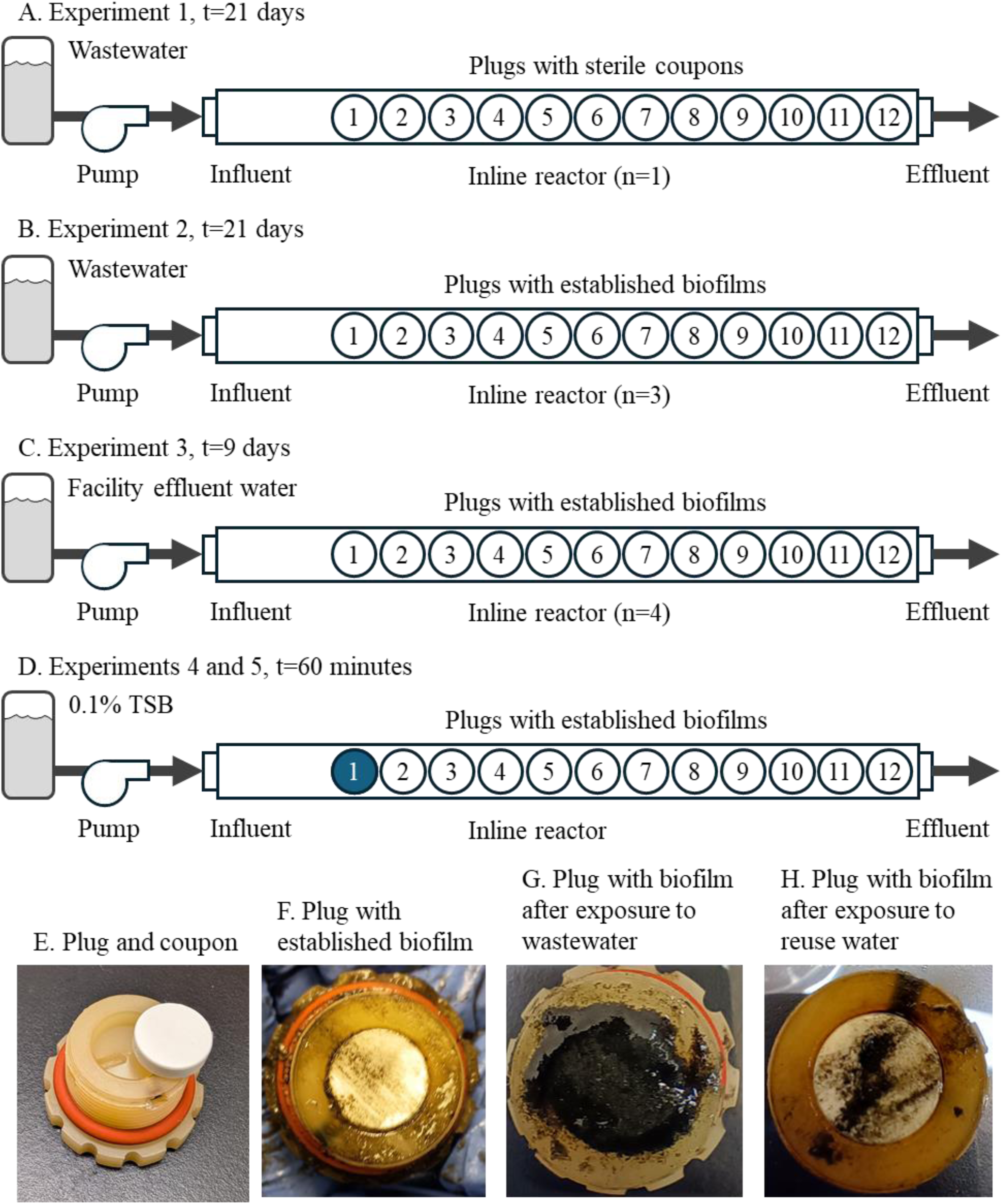
Schematic of inline biofilm reactor experiments and pictures of plugs and coupons for Experiments 1 to 5.

**Table 1.**
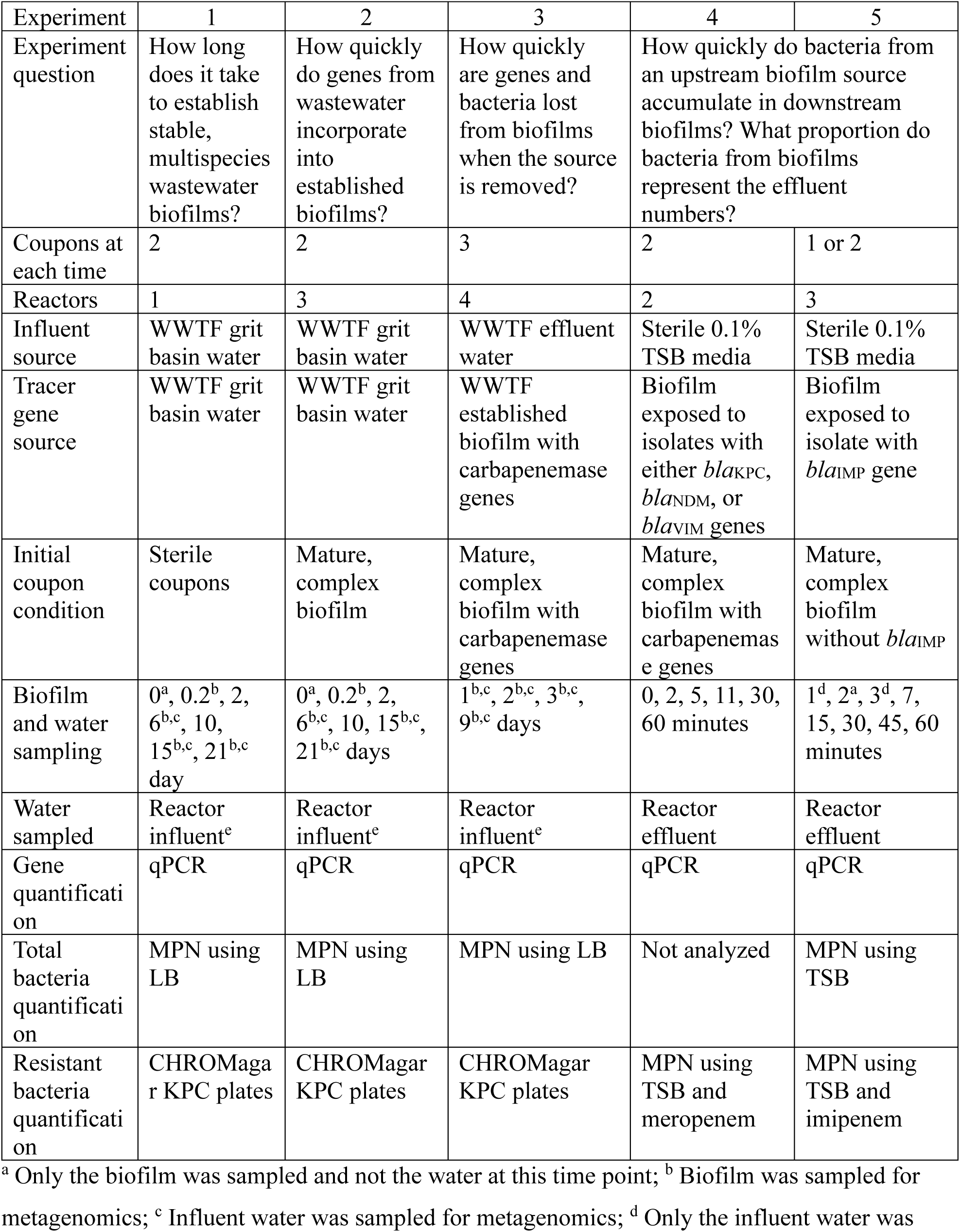
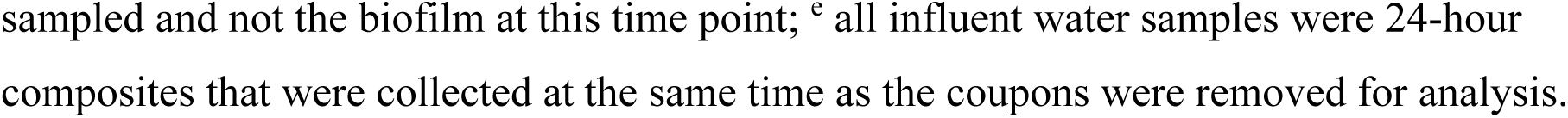
Description of the five replicated inline biofilm reactor experiments.

### 2.6 Incorporation of genes and bacteria from wastewater into an established complex biofilm

Experiment 2 was intended to model a sewer biofilm without carbapenemase genes exposed to CRO and CPOs from a colonized patient (see **Table 1** for sample collection times and analyses). Therefore, the goals for Experiment 2 were (1) to assess the accumulation of CRO and carbapenemase genes in a previously established biofilm that did not contain carbapenemase genes and (2) to monitor changes in the biofilm ecology and AMR genes over three weeks. Biofilms on PVC coupons were established by placing 44 sterile coupons in a sewer servicing a gymnasium facility for at least four weeks. The gymnasium has a daily attendance of 500 to 2000 people. After four weeks, four PVC coupon biofilms were tested for CROs and carbapenemase genes to determine background concentrations, and four PVC coupons were used for total and volatile solids of the biofilm. The remaining 36 coupons were randomly assigned to one of three replicate inline reactors and exposed to grit basin water as influent for three weeks. This experiment was run concurrently with Experiment 1. Sterile PVC coupons were used to replace any coupon removed for analysis (see **Table 1**) which were later used in Experiment 3.

### 2.7 Loss of CRO and carbapenemase genes over time after removal of selective pressure

In experiment 3, the loss of CRO and carbapenemase genes from a biofilm was evaluated to simulate the effect of a previously colonized or infected patient no longer shedding CROs or carbapenemase genes into the sewer (**Table 1**). Here, the reactor influent water in experiments 1 (n = 1 reactor) and 2 (n = 3 reactors) was switched from grit chamber to facility effluent, which contained little to no carbapenemase genes, and the microbial community was observed for nine days. As the coupons had been replaced during experiment 1 and 2, these coupons had biofilms that ranged in age from 5 to 26 days. The WWTF effluent characteristics are described in **supplemental Table S6**. The coupons were collected four times in triplicate over nine days, and the influent (i.e., a 24-hour composite) was analyzed each time a coupon was collected.

### 2.8 Detachment and transport of CPOs and carbapenemase genes from established biofilms

The purpose of experiments 4 and 5 was to determine how quickly CPOs attach to established, complex biofilms and, once in biofilms, if they detach and reattach downstream under realistic shear stresses (**Table 1**). Biofilms were established on PVC coupons in the gymnasium sewer over at least four weeks prior to this study. Nucleic acids were extracted from established biofilms on the PVC coupons (n = 2 replicates for experiment 4 and n = 4 for experiment 5) after at least four weeks of exposure to determine background carbapenemase gene concentrations. After the establishment of the complex biofilms, the coupons were exposed to either *bla*_KPC_-, *bla*_VIM_-, and *bla*_NDM_-containing isolates (experiment 4), or a *bla*_IMP_-containing isolate (experiment 5). Isolates used in this study included *Klebsiella pneumoniae* or *Pseudomonas aeruginosa* containing *bla*_KPC_, *bla*_VIM_, *bla*_NDM_, or *bla*_IMP_ (**supplemental Table S2**) that were cultured in 400 mL LB media with 5 percent sheep blood and 8 mg/L meropenem, harvested by centrifugation and then the cell pellets were resuspended in 2 mL sterile 1X PBS. The complex biofilms on PVC coupons from the sewer were then immersed in a stirred suspension of *bla*_KPC_, *bla*_VIM_, and *bla*_NDM_ isolates (n = 4 coupons for experiment 4) or *bla*_IMP_ isolates (n = 6 for experiment 5) for two minutes. After immersion of the PVC coupons in the cultures, the coupons were then rinsed with 10 mL of sterile 1X PBS to remove loosely adhered *K. pneumoniae* or *P. aeruginosa* isolates from the biofilms. Two (i.e., for experiment 4) or three (i.e., for experiment 5) of these biofilms on PVC exposed to the isolates were analyzed for starting bacteria and gene concentrations. The remaining coupons with biofilms exposed to the isolates were placed in position 1 in the reactors and served as the only CPO source for the downstream biofilms (**Figure 1D**). The remaining coupons with established biofilms from the gymnasium sewer were randomly positioned in the reactors’ remaining 11 positions downstream from the *K. pneumoniae* or *P. aeruginosa* isolate exposed coupons.

The influent solution was sterile 0.1% TSB to simulate wastewater flowing at 10 L/hour representing shear stresses in the sewer line. At this flow rate, the 98 ml interval volume of the reactor was replaced 102 times over the one hour of the experiment. Over 1 hour of flow, duplicate coupons from each reactor were removed from random locations and analyzed as described in **Table 1** including culture and qPCR-based analysis of CRO and carbapenemase genes. Removed coupons were replaced with sterile PVC coupons. In experiment 5, the lowest MPN dilutions that were positive for growth were extracted and tested for *bla*_IMP_ genes by qPCR in a nested qPCR approach to lower the sensitivity of the analytical method. Six to seven effluent samples (**Table 1**) were collected over the experimental period and analyzed for total and resistant bacteria and genes. A mass balance approach was used to assess the transfer of CPO and carbapenemase genes from the position 1 coupon in the reactor to the effluent (see supplemental information for mass balance calculations).

## 3.0 RESULTS

### 3.1 Wastewater and facility effluent carbapenemase genes, bacteria and meropenem

Meropenem was not detected in either the grit basin wastewater or the WWTF effluent above the minimum detection limit of 0.01 µg/mL, which is well below the minimum inhibitory concentration for carbapenem resistance of 1 to 8 µg/mL [28]. Grit basin wastewater showed little variation over the 21-day study period with 7.1 ± 0.3 log 16S rRNA GC/mL (average ± standard deviation) (**Figure 2**). Total bacterial concentrations ranged from 7.0 to 7.7 log MPN/mL, while CROs were 5.6 to 6.4 log CFU/mL (**Figure 2**). The carbapenemase genes were 2 to 4 orders of magnitude less than the 16S rRNA GC/mL and averaged 3.5 ± 0.14, 5.7 ± 0.24, 4.5 ± 0.22, 4.1 ± 0.23, 4.4 ± 0.16 log GC/mL for *bla*_KPC_, *bla*_OXA-48-like_, *bla*_VIM_, *bla*_NDM_, and *bla*_IMP_, respectively. Grit chamber water parameter values, temperature, pH, carbonaceous biochemical oxygen demand, total suspended solids, Total Kjeldahl Nitrogen, total phosphorus, and conductivity for the grit basin water are shown in **supplemental Table S5**.

**Figure 2.**
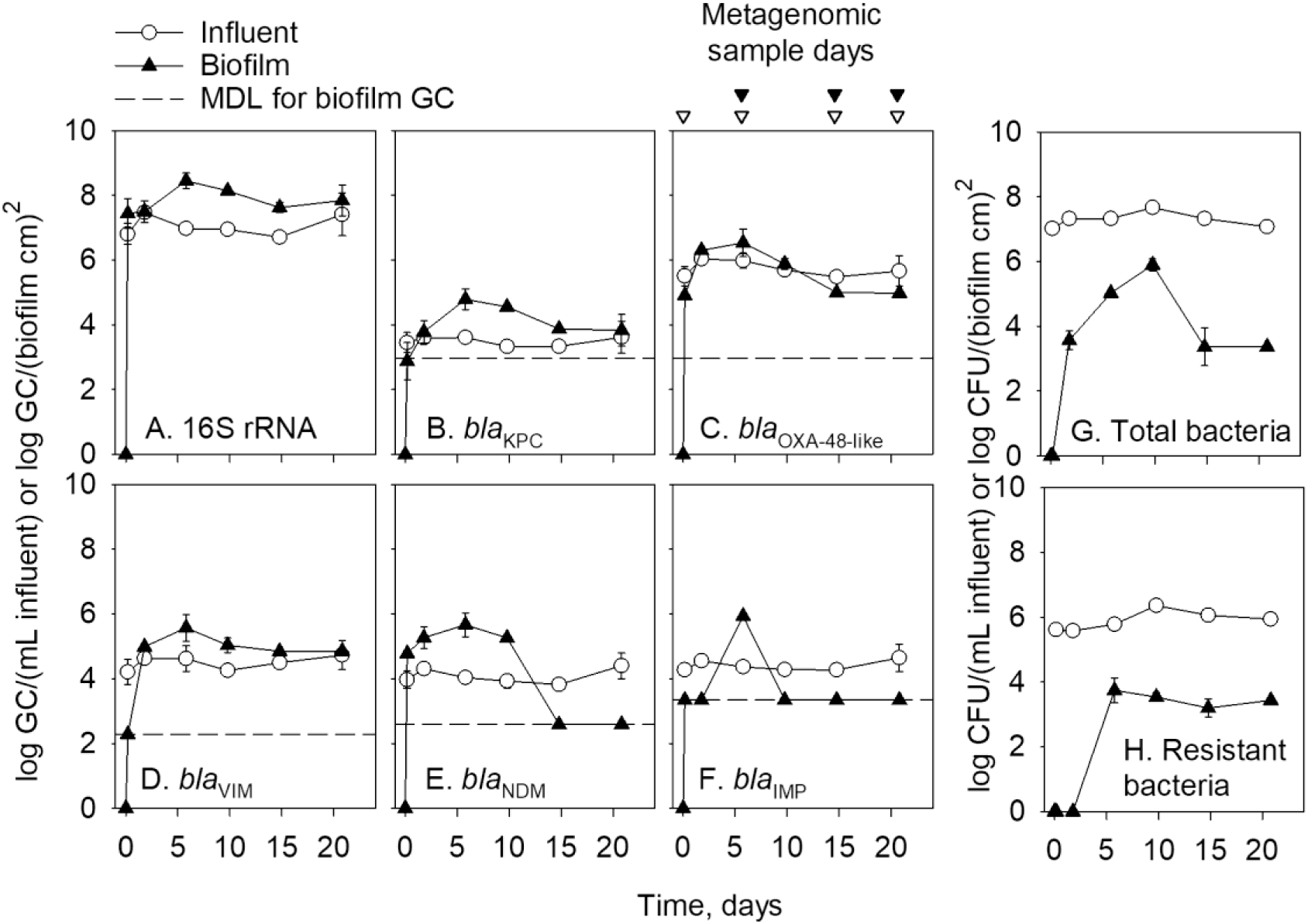
Experiment 1 gene concentrations (in log gene copies (GC) in plots A through F) and total and carbapenem-resistant organisms (log colony forming units (CFU) in plots G and H) in biofilms and influent (grit basin water) in one initially sterile reactor. Each data point is the average and standard deviation of two coupons or three influent samples. Not all error bars are visible due to small standard deviations. Coupons that were initially sterile (i.e., day 0) were plotted as log 0.1 log GC/(biofilm cm)^2^ for visualization of the temporal relationship and do not represent a measured value. Non-detect biofilm samples were plotted at the method detection limit (MDL) [22] for any of the target carbapenemase genes. Inverted triangles above plot C indicate the samples evaluated for metagenomics of the biofilms (open) or influent (closed).

Wastewater treatment facility effluent contained significantly lower 16S rRNA and CROs compared to the grit chamber water. The facility effluent averaged 2.1 ± 0.3 log GC/mL for 16S rRNA gene (**Figure 3**). *bla*_KPC_ was detected in three of four samples, averaging -0.8 ± 0.2 log *bla*_KPC_ GC/mL. Both *bla*_OXA-48-like_ and *bla*_VIM_ were detected in only one of the four samples, and *bla*_NDM_ and *bla*_IMP_ were not detected at all. The concentration of total bacteria in the effluent was less than 0 log MPN/mL after one day, then increased to 4.4 ± 0.4 log MPN/mL (average ± standard deviation) (**Figure 3**). CRO concentrations had a similar trend, with less than 0 log CFU/mL for the first day and an average of 4.2 ± 0.3 log CFU/mL for the rest of the experiment. Facility effluent parameter values are found in **supplemental Table S6**.

**Figure 3.**
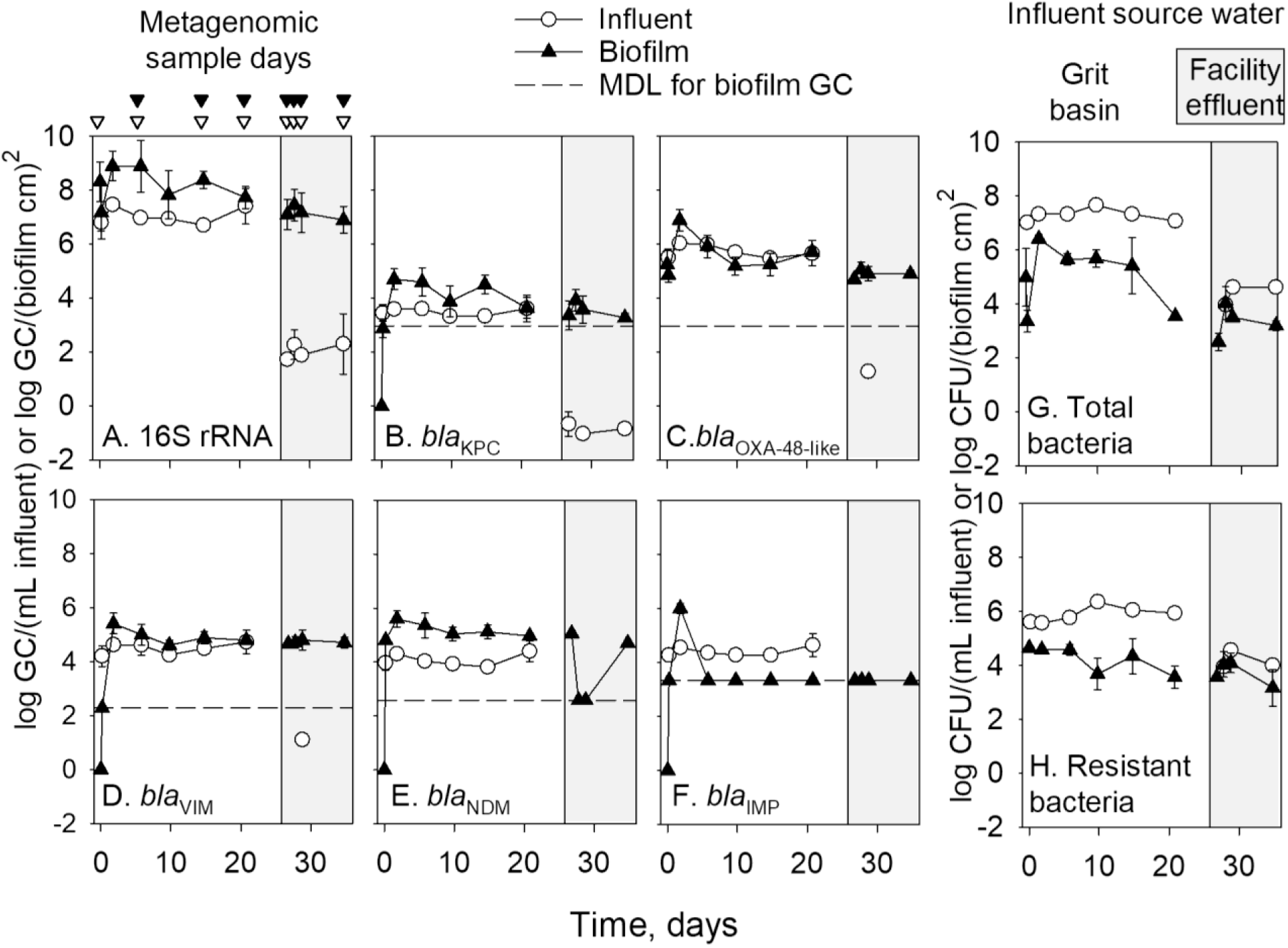
Experiment 2 and 3 gene concentrations (log gene copies (GC) in plots A through F) and total and carbapenem-resistant organisms (log CFU in plots G and H) in biofilms and influent (grit basin water for Experiment 2 or treated effluent water for Experiment 3). Experiment 2 included three inline biofilm reactors with established biofilms on coupons and ran for 26 days using grit basin water as the influent. For Experiment 2 each data point is the average and standard deviation of 6 data points (n = 2 coupons from each of three reactors) except time 0. Experiment 3 is indicated by a gray background and included four inline biofilm reactors with biofilm on coupons grown for 5 to 26 days and then exposed to effluent water as the influent to the reactors. For Experiment 3, each data point is the average and standard deviation of 12 total coupons (n = 3 coupons from each of four reactors). Coupons that were non-detect for target carbapenemase genes were plotted as log 0.1 log GC/(biofilm cm)^2^ for visualization of the temporal relationship and do not represent a measured value. Non-detect biofilm samples were plotted at the method detection limit (MDL) [22] for any target carbapenemase genes. Inverted triangles above plot A indicate the samples evaluated for metagenomics of the biofilms (open) or influent (closed).

### 3.2 Total and volatile solids of biofilms

Total and volatile solids in biofilms from the gymnasium sewer during establishment of the complex biofilms were 1.75 ± 0.32 mg/cm^2^ and 0.64 ± 0.00 mg/cm^2^, respectively, as measured in four background coupons (i.e., 37% volatile solids, see **Figure 1F**). Some biofilm coupons from the WWTF (Experiments 1 and 2) were covered in black grit, likely from the influent grit wastewater (**Figure 1G**), influenced by ferric chloride added for hydrogen sulfide control in the collection system. Total solids from the black grit biofilms were 0.53 ± 0.008 mg (based on four measurements) while volatile solids were 0.03 ± 0.002 mg, indicating the black grit observed on the PVC coupons was not organic and likely an iron hydroxide.

### 3.3 Time required to establish stable, multispecies biofilm from wastewater

A stable biofilm was established within one to two weeks on previously sterile PVC coupons exposed to grit basin influent in experiment 1. Within two days of exposure to wastewater, carbapenemase and 16S rRNA genes increased between 3 and 7 log GC/cm^2^. Most genes on the PVC coupons reached maximum concentrations by day six (**Figure 2A** to **2F**). After six days, some carbapenemase gene concentrations decreased in the biofilms, but typically by less than 1 log GC/cm^2^, indicating a relatively stable community. Similarly, total bacterial concentrations increased over 10 days to 6 log MPN/cm^2^ before decreasing to a steady 3.4 log MPN/cm^2^ for the remainder of the experiment (**Figure 2G**). CROs reached maximum concentration at day six and thereafter averaged 3.5 ± 0.2 log CFU/cm^2^.

Additional evidence of the biofilm stability was evaluated using the overall taxonomic profile of the community and composition of AMR genes with metagenomic sequencing. The operational taxonomic unit (OTU) compositions of the biofilm metagenomes on the initially sterile coupon (reactor 4; experiment 1) were at first distinct from those of the other reactors with mature biofilms and closer to that of the influent water (**Figure 4**). On days 15 and 21, the OTU compositions of the newly developing biofilm in reactor 4 became more similar to, but still distinct from, those of the mature biofilms. The number of OTUs in these biofilm metagenomes were generally constant after day 6 (**supplemental Figure S1)**.

**Figure 4.**
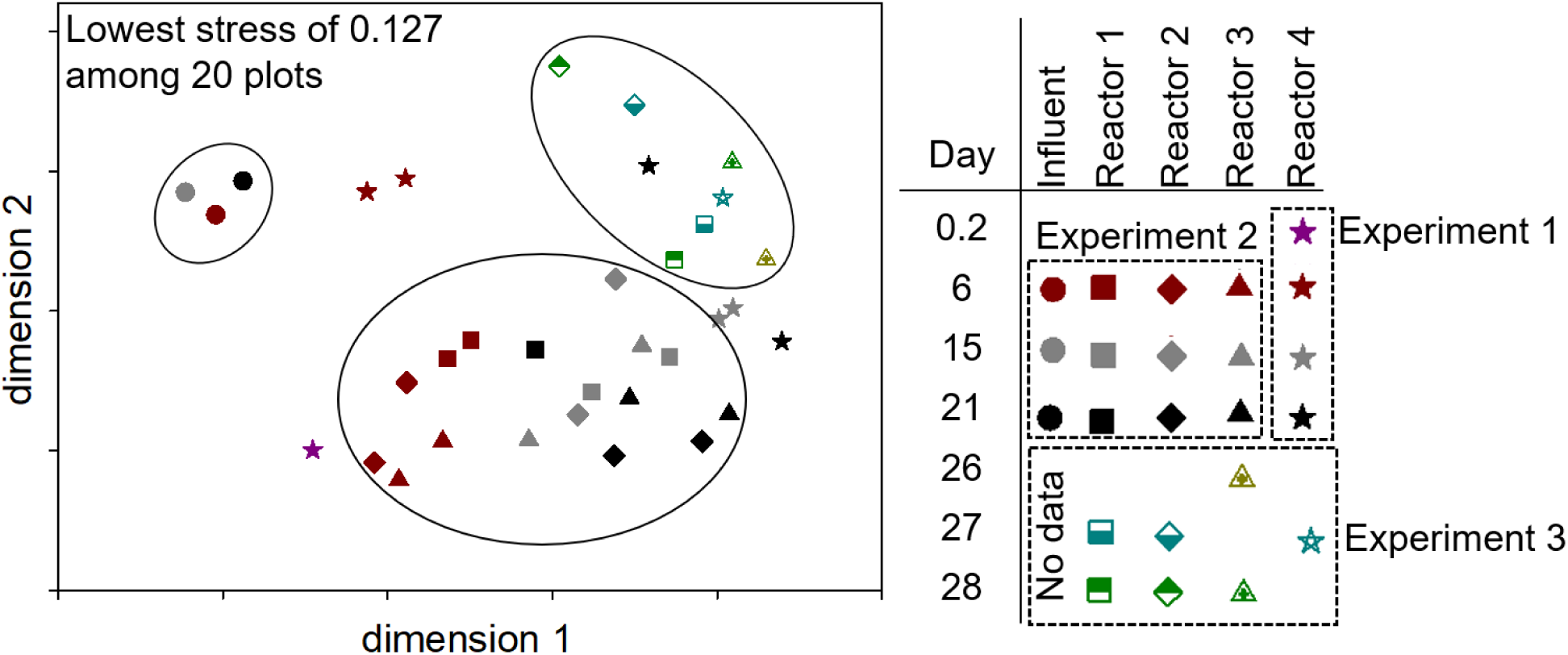
Non-metric multidimensional scaling (NMDS) analysis of influent and biofilm bacteria OTUs in metagenomes over time in experiments 1, 2, and 3 (indicated by dashed boxes in the legend). Reactors 1 to 3 (Experiment 2 replicate reactors) started with coupons with mature biofilms; Reactor 4 (Experiment 1) started with sterile coupons. After 25 days the influent for all four reactors was switched to effluent water for Experiment 3. Circled data points in the NMDS plot are groups of influent samples for Experiments 1 and 2, established biofilm samples of Experiment 2, and biofilm samples of Experiment 3. “No data” indicates the samples were subjected to metagenomics sequencing, but due to low DNA input, very few sequences were obtained.

The most abundant AMR gene categories were macrolide, beta-lactam, macrolide/streptogramin, lincosamide/macrolide/streptogramin, tetracycline, aminoglycoside, and sulfonamide (**Figure 6**). The most abundant individual genes in most biofilm metagenomes were the macrolide genes *mph*(E) and *msr*(E) and the lincosamide/macrolide/streptogramin gene *erm*(B). These genes are especially abundant at the earliest time points and are greatly reduced in their relative contribution in days 15-21 of the newly developing biofilm (**supplemental Figure S2**).

Most carbapenemase genes targeted for qPCR were not detected in any of the metagenomes (i.e., *bla*_KPC_, *bla*_VIM_, *bla*_NDM_, and *bla*_IMP_). The *bla*_OXA_ family is abundant in most biofilm metagenomes, but *bla*_OXA-48_ does not appear as a specific annotation. The most abundant *bla*_OXA_ sequence is not identified with any more specificity by AMRfinder, but the closest blast hit (99 percent amino acid identities over 84 percent of the predicted gene) is a *Pseudomonas* OXA-2 family class D beta-lactamase OXA-544. This *bla*_OXA_ gene was highly abundant on the sterile coupon on day 0.2 but decreased in relative abundance at later time points (supplemental data files).

### 3.4 Incorporation of genes and bacteria into mature multispecies biofilms from wastewater

CRO and carbapenemase-gene concentrations in biofilms stabilized within two days when established biofilms were exposed to grit chamber wastewater, as shown in experiment 2 (**Figure 3**). The 16S rRNA gene concentration in the biofilm was initially 8.3 GC/cm^2^ and decreased by 1.2 log GC/cm^2^ after four hours of exposure to the grit basin water. Concentrations then rebounded to an average of 8.3 log GC/cm^2^ for the rest of the experiment (**Figure 3A)**. The target carbapenemase genes in the biofilms increased quickly within the first four hours of exposure to the high concentration grit chamber water from non-detect in the biofilm to 4.0 ± 0.7, 4.5 ± 1.1, 5.2 ± 0.3, and 3.8 ± 1.1 log GC/cm^2^ for *bla*_KPC_, *bla*_VIM_, *bla*_NDM_, and *bla*_IMP_, respectively, within two days of exposure.

As with the 16S rRNA gene concentrations, total bacterial concentrations in the biofilm decreased briefly after four hours but rebounded quickly after two days to their highest level at 6.4 log MPN/cm^2^ (**Figure 3G)**. Total bacterial concentrations trended lower over the next 19 days and ended at 3.6 log MPN/cm^2^, likely due to increased shear stress associated with particulate matter from the grit chamber. CRE started at 4.6 log CFU/cm^2^ and decreased by 1.0 log CFU/cm^2^ during the experiment when grit-chamber water was used as the influent to the reactor (**Figure 3H**). At the start of the experiment with grit-chamber influent, the influent contained 5 ± 2% meropenem-resistant bacteria. In the biofilm, 47% of the total bacteria were also meropenem resistant, but this decreased to 25 ± 45% by the end of 21 days of observation.

The Shannon diversity index of the metagenomic OTUs and a number of distinct AMR genes were relatively stable from day 6 to day 21 in the established biofilms (**supplemental Figure S1** and **S2**). The community composition of metagenomic OTUs and AMR genes in the three reactors for experiment 2 (“reactor 1” to “reactor 3” from Day 0.2 to Day 21 in **Figures 4** and **5**) shifted over time, with the community composition at Day 6 being closer to the composition of influent water than later time points. This trend over time for OTU composition (**Figure 4**) was similar but less pronounced for the composition of AMR genes (**Figure 5**).

**Figure 5.**
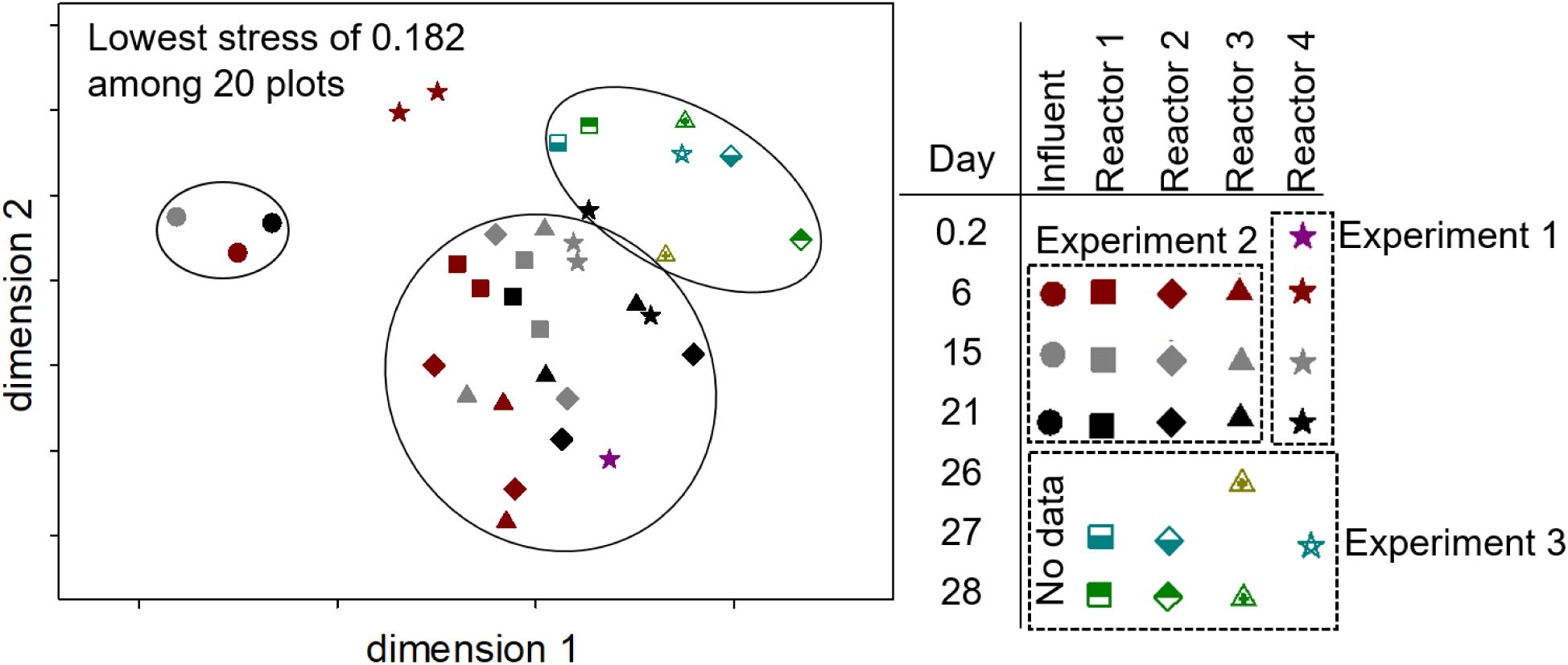
Non-metric multidimensional scaling (NMDS) of antimicrobial resistance genes in the metagenomes of the influent and in biofilms over time in experiments 1, 2, and 3 (indicated by dashed boxes in the legend). Reactors 1 to 3 (Experiment 2 replicate reactors) were started with mature biofilms; Reactor 4 (Experiment 1) was started with sterile coupons. After 25 days the influent for all four reactors was switched to effluent water for Experiment 3. Circled data points in the NMDS plot are groups of influent samples for Experiments 1 and 2, established biofilm samples of Experiment 2, and biofilm samples of Experiment 3. “No data” indicates the samples were subjected to metagenomics sequencing, but due to low DNA input, very few sequences were obtained.

**Figure 6.**
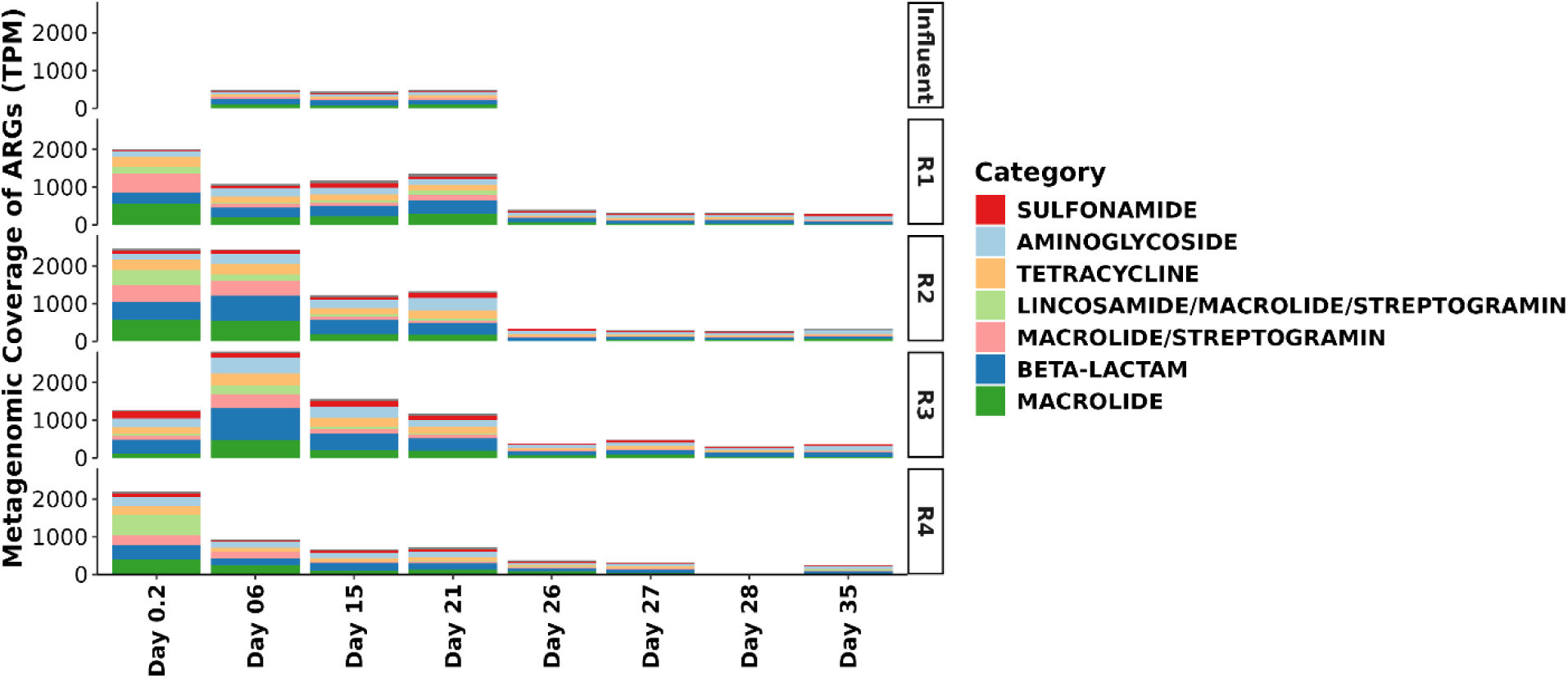
Metagenomic coverage of antimicrobial-resistance genes (ARGs) by day for each reactor. R1 through R3 represent triplicate reactors in Experiment 2 and 3. R4 was the single reactor in Experiment 1. Influent was the grit chamber wastewater for Days 0 to 21 and effluent water for Days 26 to 35. Due to the low DNA available from the effluent water, no metagenomic data are available for those samples (“Influent” for Days 26-35).

The distribution of AMR genes in the mature biofilms was similar to that of the newly developing biofilms, with common gene categories and the same dominant genes (e.g. *mph*(E), *msr*(E), *erm*(B), and *bla*_OXA_). However, the relative abundances of AMR genes were generally much higher in the mature biofilms (**Figure 6**). Sequences annotated as *bla*_OXA_, *bla*_OXA-2_, and *bla*_MCA_ remained at much higher abundances on days 15-21 compared to the newly developing biofilm. Like the metagenomes of the developing biofilms (experiment 1), none of the target genes for qPCR (*bla*_KPC_, *bla*_VIM_, *bla*_NDM_, and *bla*_IMP_) were detected in the mature biofilms. *bla*_OXA-48_ was not detected, but 58 other beta-lactamase genes were identified.

### 3.5 Loss of carbapenemase genes and CRO from biofilms

After switching from grit chamber water to WWTF effluent water as influent in the replicate biofilm reactors, we observed persistence of target carbapenemase genes and resistant bacteria in the biofilm over eight days (**Figure 3**). The facility effluent water contained significantly lower 16S rRNA, *bla*_KPC,_ and BOD than the grit chamber water (ANOVA, *P*<0.001, n = 6 for grit chamber water and n = 4 for effluent water). Further, *bla*_OXA-48-like_ and *bla*_VIM_ were only detected once over eight days in the facility effluent, and *bla*_IMP_ was not detected. While 16S rRNA, *bla*_KPC_, and *bla*_OXA-48-like_, decreased over the eight days following the switch to facility effluent as the reactor input, this change was typically less than 1 log decrease in GC/cm^2^ (grey areas of **Figure 3A**, **3B,** and **3C**). Total bacteria and CRE persisted in the biofilm over eight days, even after switching to a low BOD influent. The total bacteria initially increased from 2.6 MPN/cm^2^ to 4.0 MPN/cm^2^ but then decreased to 3.4 MPN/cm^2^ (**Figure 3G**), with resistant bacteria following a similar pattern (**Figure 3H**). These results suggest that target carbapenemase genes and total and resistant bacteria take longer than eight days to respond to changes in influent water characteristics under realistic shear conditions. While target carbapenemase gene copies and culture counts suggest minimal changes in the biofilm with a switch from grit chamber wastewater to facility effluent (**Figure 3** and **Figure 6**), metagenomic coverage of all AMR genes declined significantly during this period (**Figure 5**, days 26 to 35). Furthermore, the samples exposed to effluent water had distinct compositions of OTUs and AMR genes, with multiple species and genes changing their abundance compared to the grit chamber samples (**Figures 4** and **5**). While most AMR genes decreased in relative abundance after the switch to facility effluent water, a few (e.g., *aadA7* and *bla*_OXA-10_) showed moderate increases. This persistence mirrors observations from healthcare plumbing systems, where carbapenem-resistant organisms can remain despite removal of antibiotic selective pressure, contributing to recurrent exposure risks for patients and complicating remediation efforts [24, 26].

### 3.6 Release from and reattachment of CROs and carbapenemase genes in sewer biofilms

Biofilms downstream of a single biofilm coupon containing high concentrations of *P. aeruginosa* with *bla*_KPC-5_, *K. pneumoniae* with *bla*_VIM-27_, and *K. pneumoniae* with *bla*_NDM-1_ remained within 0.5 log GC/cm^2^ of the starting concentration over 60 minutes of exposure to high shear flow (experiment 4, **Table 2**, **Figure 7**). Even though the biofilms were not exposed to *bla*_OXA-48-like_-containing cultures, measurable concentrations were found in the established coupons, likely originating from the wastewater used to establish the original biofilm. Once flow started, concentrations of 16S rRNA genes immediately increased in the effluent to 7.3 log GC/mL, then decreased to 2.8 log GC/mL by the end of the experiment (**Figure 7A**), suggesting significant sloughing of biofilm from the PVC coupons. Trends in *bla*_OXA-48-like_, *bla*_VIM_, and *bla*_NDM_ acted similarly to 16S rRNA in the effluent. After the initial increase to 3.2 log GC/mL, *bla*_KPC_ concentrations decreased for 30 minutes, then increased to 2.3 log GC/mL after 60 minutes (**Figure 7B**). Meropenem-resistant bacteria in the biofilm increased over the first five minutes to 7.4 log CFU/cm^2^, after which the concentration remained relatively unchanged (**Figure 7F**). Due to the presence of the carbapenemase genes in the biofilm prior to exposure to the culture, we were unable to perform a mass balance on them in this study. However, the presence of bacteria and genes in the effluent suggests that carbapenemase genes are released into the flowing water above the biofilms and the genes persist in mature biofilms over 60 minutes of high shear flow.

**Figure 7.**
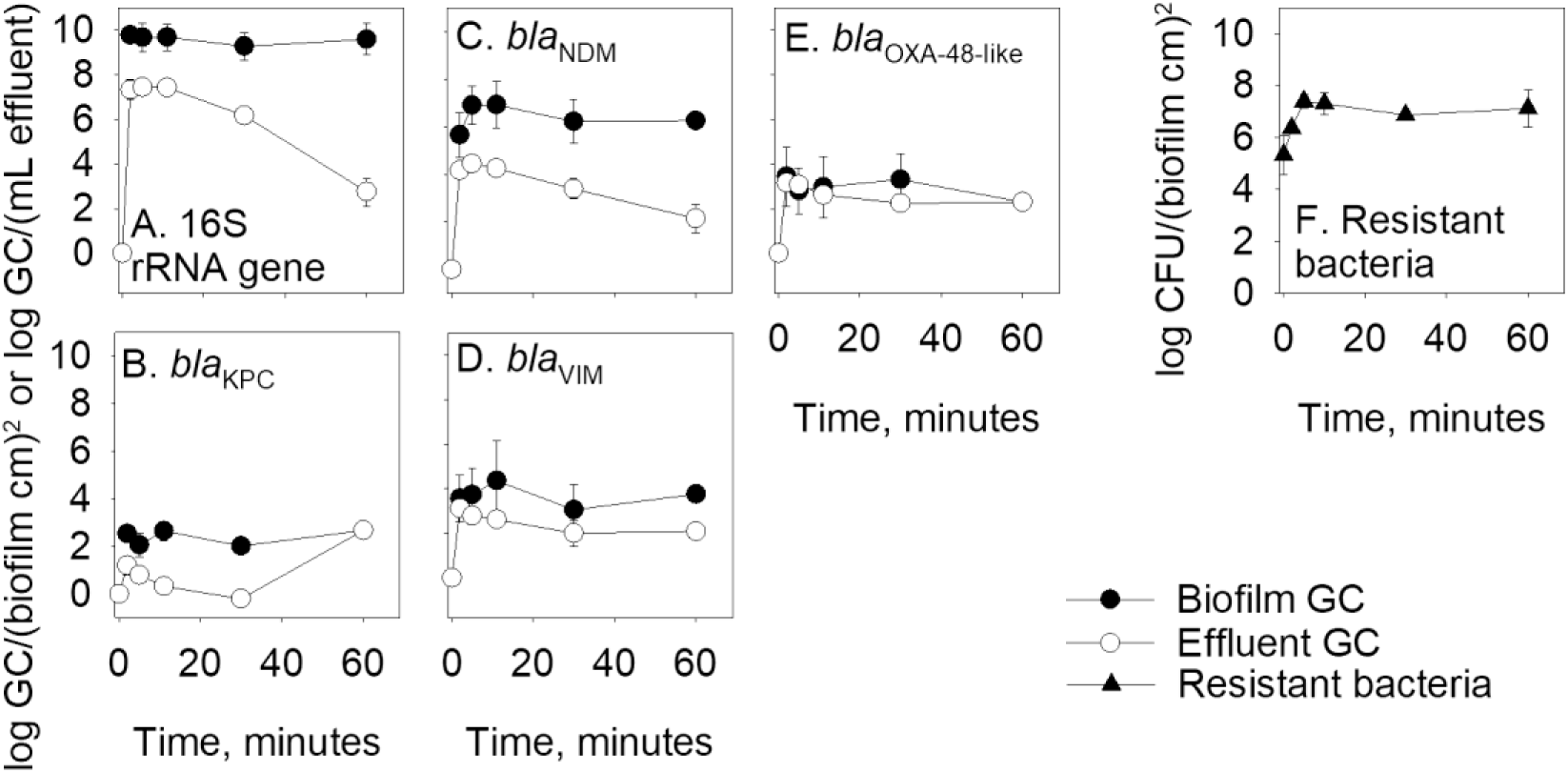
Inline biofilm reactors run in duplicate for 60 minutes with *bla*_KPC_-, *bla*_VIM_-, and *bla*_NDM_-containing bacteria added to mature biofilms in position 1 in the reactor (Figure 1, Experiment 4). Results represent the average and standard deviation of four samples, except for the point at 60 minutes, which was based on two samples. Non-detects were replaced with one-half the detection limits in the average calculations.

**Table 2.**
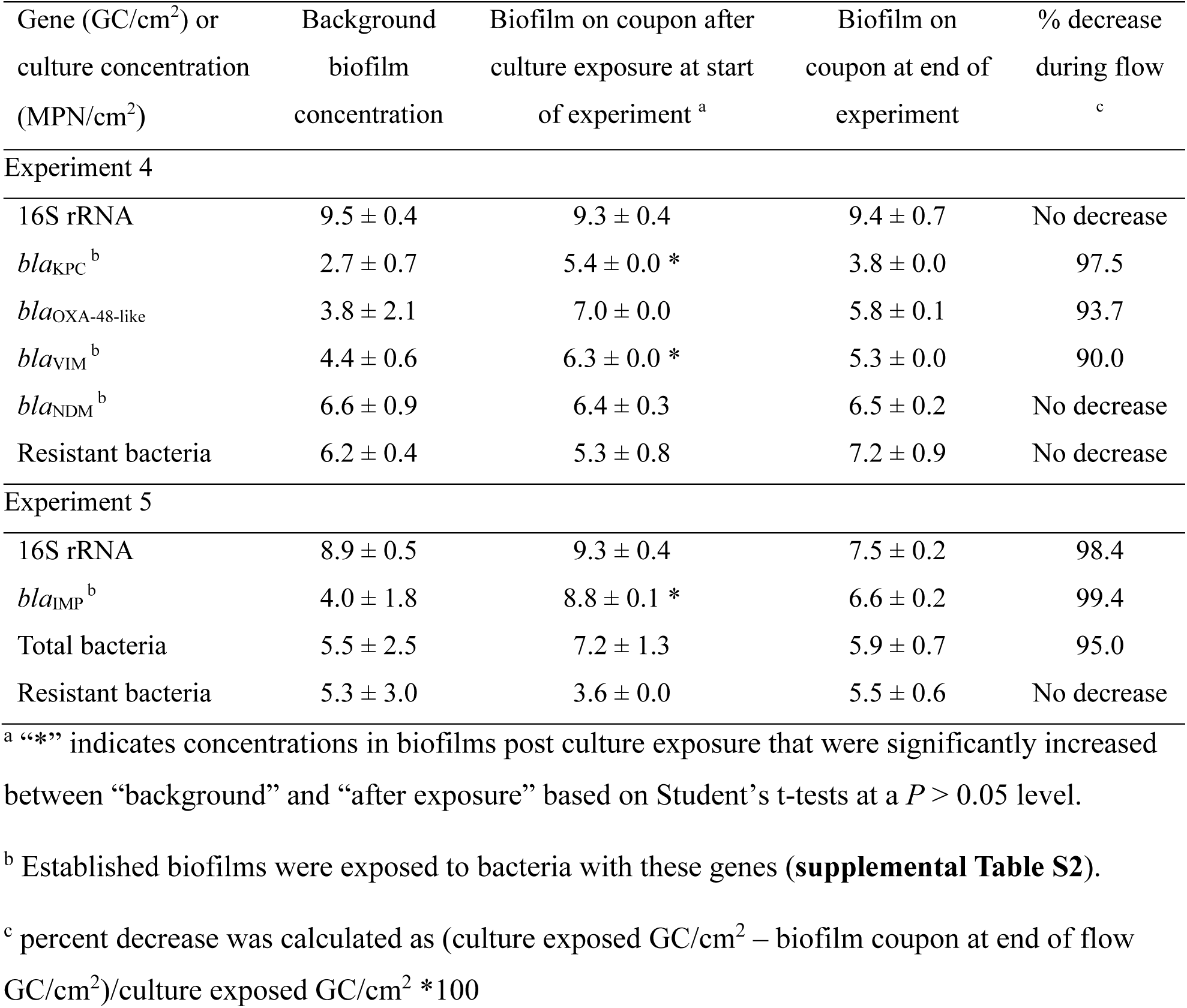
Biofilm concentrations after exposure to carbapenemase-producing organisms, *Pseudomonas aeruginosa* or *Klebsiella pneumoniae*, in Experiments 4 and 5 in average ± standard deviation of log gene copies (GC)/cm^2^ or log most probable number (MPN)/cm^2^.

When this experiment was repeated (i.e., experiment 5) with a *bla*_IMP_-containing *P. aeruginosa* culture added to established biofilms, the mass balance between the biofilm and the effluent was estimable (**Table 3**), and we were able to evaluate the transfer of *bla*_IMP_ to downstream biofilms (**Figure 8**). The *bla*_IMP_-containing *P. aeruginosa* were translocated from the first PVC coupon to downstream coupons, but at very low concentrations over 60 minutes. Specifically, culturing scraped biofilm in TSB media containing imipenem over 24 hours revealed that imipenem-resistant bacterial cultures contained *bla*_IMP_ detected by qPCR. However, *bla*_IMP_ was not amplifiable from the scraped biofilm without prior culturing, indicating low concentrations of CPO are transported downgradient and adhere to mature biofilms (**Figure 8C**). The *bla*_IMP_ gene was detectable in the reactor effluent for the first 10 minutes (**Figure 8D**).

**Figure 8.**
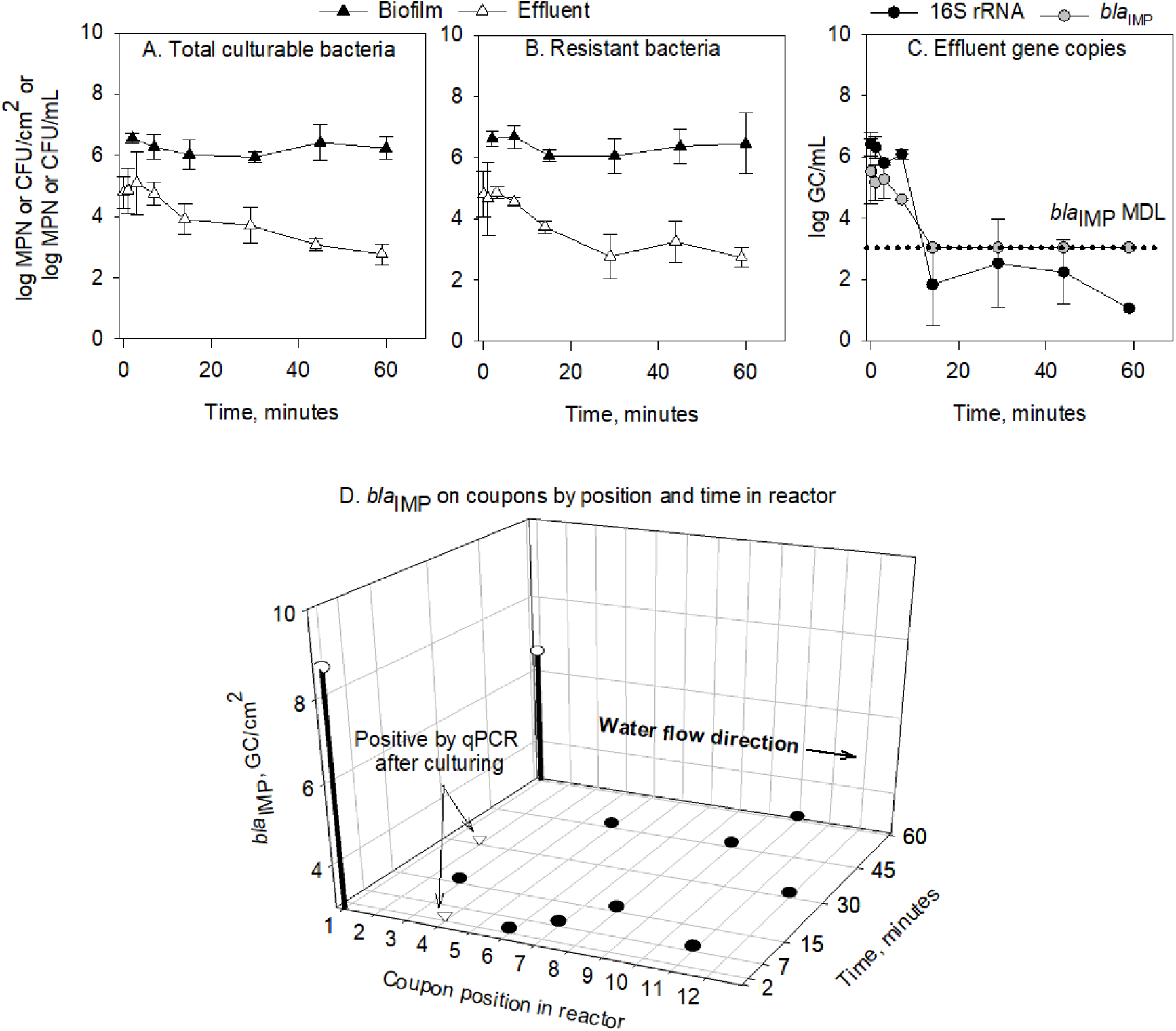
Inline biofilm reactors run in triplicate for 60 minutes with *bla*_IMP_-containing bacteria as a source in position 1 (Figure 1, Experiment 5). A. Total bacteria in biofilm and effluent. MPN = most probable number; CFU = colony-forming units. B. Imipenem-resistant bacteria in the biofilm and effluent. C. 16S rRNA and *bla*_IMP_ genes in effluent. GC = gene copies. D. *bla*_IMP_ genes in the biofilms. Arrows and upside-down triangles indicate biofilm samples that were positive for *bla*_IMP_ by qPCR assay after enrichment in media with meropenem. Black-filled circles indicate biofilm samples that were tested, but were non-detect for *bla*_IMP_. Data points represent six reactor biofilm samples and three laboratory splits for effluent samples. Non-detects were replaced with one-half the detection limits [22] in the average calculations.

**Table 3.** Mass balance of genes (total gene copies) or culturable bacteria (total most probable numbers) between biofilms and effluent, pre- and post-flow in Experiment 5.

|  | 16S rRNA<br>genes | *bla*<sub>IMP</sub><br>genes | Total<br>bacteria | Resistant<br>bacteria |
| --- | --- | --- | --- | --- |
| <u>Start</u> |  |  |  |  |
| Background <sup>a</sup> | 9.6 ± 0.5 | 6.6 ± 1.8 | 7.4 ± 2.5 | 7.8 ± 3.0 |
| Start <sup>b</sup> | 9.9 ± 0.4 | 9.3 ± 0.1 | 8.5 ± 1.3 | 4.2 ± 0.0 |
| <u>End</u> |  |  |  |  |
| Coupon <sup>c</sup> | 8.0 ± 0.3 | 7.2 ± 0.2 | 6.8 ± 0.7 | 6.3 ± 0.6 |
| Effluent <sup>d</sup> | 9.5 ± 0.1 | 8.8 ± 0.3 | 8.7 ± 0.3 | 8.4 ± 0.3 |
| Recovery | 40% | 36% | 150% | 110% <sup>e</sup> |

“Start” values are the initial values (gene copies or most probable number) in the biofilm in the reactor. “End” values are the final values after the experiment. “Recovery” values are the percent of gene copies that are accounted for in the biofilm and effluent at the end of the experiment.
<sup>a</sup> average of 4 biofilms
<sup>b</sup> average of 3 biofilms
<sup>c</sup> average of 3 biofilms from 3 reactors
<sup>d</sup> total of flow-weighted values
<sup>e</sup> This calculation used the background concentration since the starting resistant bacteria was lower.

Compared to the large decrease in *bla*_IMP_, 16S rRNA gene concentrations in the reactor biofilms only decreased from background concentrations by 0.8 log GC/cm^2^ and similarly, total bacteria in biofilms varied by only ± 0.3 log MPN/cm^2^ (**Figure 8C** and **Figure 8A**). Bacteria in the effluent decreased from 5.1 log MPN/mL to 2.8 log MPN/mL over the experiment. Resistant bacterial concentrations in the biofilm and effluent were not significantly different from those of total bacteria (**Figure 8B** and **Figure 8D**, ANOVA, P = 0.379), indicating that most culturable bacteria in the effluent are resistant to at least 8 mg/L imipenem.

Mass balances for the 16S rRNA and *bla*_IMP_ genes in Experiment 5 showed 40 and 36 percent recovery, respectively (**Table 3**). In each case, there were more genes found in the effluent (9.5 log GC for 16S rRNA and 8.8 log GC for *bla*_IMP_) than in the biofilms (8.0 log GC 16S rRNA and 7.2 log GC *bla*_IMP_). The decrease in gene mass from start to end may be due to bacterial adhesion to the reactor walls or to errors in measuring the effluent. For bacterial counts, total and resistant bacteria at the end of the experiment were 150 and 110 percent of the initial numbers, respectively.

## 4.0 DISCUSSION

This study demonstrates that sewer biofilms act as dynamic reservoirs of CRO and CP-CRE, with exchange of bacteria and resistance genes occurring across multiple time scales. Uptake of resistant organisms and genes into previously established complex biofilms occurs within minutes to hours, while incorporation into downstream biofilms reaches quasi-steady concentrations within 1-2 days. These rapid kinetics suggest that mass transfer of bacteria and genes into sewer biofilms is not diffusion-limited under the tested conditions, but instead controlled by hydrodynamic delivery and surface attachment processes. This rapid uptake of CRO and carbapenemase-producing CRE into sewer biofilms is consistent with high-frequency exposure events in building-scale sewers from toilet flushing, where short duration pulses of elevated microbial concentrations repeatedly contact biofilm surfaces. Under these conditions, biofilms may integrate transient inputs into a time-averaged reservoir of microorganisms and genes. Other studies have also reported stable biofilms after 10 days [29] or within two weeks [30]. Sulfide production, a bulk measurement of biofilm formation, in the biofilm stabilized after two weeks [30] while biofilm thickness and density stabilized within 20-30 days under different shear stresses [31]. While concentrations of bacteria and genes stabilized within days, metagenomic results herein indicate that the biofilm continued to evolve over the first two weeks. Other metagenomic analyses reported in the literature showed that the biofilm bacterial species changed over 13 weeks to one year [30, 32]. This study and others report that biofilms form quickly and that bacteria and gene concentrations stabilize within days to weeks, but biofilm community composition continues to change over longer time periods.

Despite rapid uptake of CRO and CP CRE genes, removal of selection pressure by switching to water with few bacteria, low biochemical oxygen demand and low antibiotic concentrations (i.e., non-detectable meropenem), resulted in minimal loss of resistance genes and bacteria over 8 days (i.e., < 1 log reduction) under the same shear stress. These results are in contrast with those of Stoodley et al. [36], which found a 10-fold change in biofilm bacterial species concentrations within a day after increasing and then decreasing glucose concentrations by a factor of 10. The persistence of genes in this study suggests that sewer biofilms may provide a “memory”, retaining bacteria and genes from wastewater long after the upstream inputs have ceased. Importantly, not all resistance markers behaved similarly in this study, highlighting that microbial and genetic traits influence attachment and downstream movement. Further, while qPCR and culture-based concentrations remained relatively stable, metagenomic analyses revealed continued shifts in microbial community composition over weeks. In field samples, metagenomic analyses of wastewater show temporal shifts in Bray-Curtis Dissimilarities over weeks to months of sampling [33, 34]. These shifts are likely due to changing conditions [35]. Therefore, the decoupling observed herein suggests that bulk gene concentrations alone (e.g., 16S rRNA or specific resistance genes) are insufficient to characterize biofilm dynamics, and that underlying community restructuring may occur without large changes in absolute abundance.

The central implication from these findings is that sewer biofilms may decouple wastewater signals from real-time human shedding of genes and bacteria of interest from an epidemiology perspective. Rather than sewer systems acting as passive conduits, sewer biofilms may rapidly attenuate peaks through immediate adsorption, store microorganisms and genes, and gradually release them over time, producing delayed and dampened signals. This behavior transforms wastewater signals from discrete pulses to broadened, time-integrated profiles, complicating the interpretation of wastewater surveillance results. As a result, composite samples may over-represent historical contributions, and observed persistence of resistance genes may reflect biofilm reservoirs rather than ongoing infections. Other authors have recommended sampling of sewer biofilms to assess their impact on wastewater [7, 8]. Along with biological stability, dynamic wastewater flow, physical and chemical properties, and sediments, understanding biofilm interactions can help elucidate bacterial and viral transport in sewers [8]. Biofilms can act as reservoirs of AMR and contribute to its downstream spread [7]. This apparent biofilm “memory” has important clinical implications, as wastewater signals may lag true patient-level transmission dynamics. Sewer biofilms are also important due to their influence on chemical biomarker stability [36]. These findings challenge a core assumption of WBE that measured concentrations reflect near real-time upstream human inputs. In healthcare settings, this could delay recognition of outbreak resolution or obscure the effectiveness of infection control interventions [21]. Results herein highlight the need to incorporate biofilm-mediated storage and release into WBE models. This apparent biofilm “memory” has important clinical implications, as wastewater signals may lag true patient-level transmission dynamics.

While this study has shown that sewer biofilms are influenced by changing wastewater flowing over them, several additional points warrant investigation in further studies. First, sewers are made of different materials beyond PVC including cast iron, concrete, vitrified clay, and fiber conduit pipe (also known as Orangeburg), and these should be tested to model biofilm formation and exchange. Second, wastewater flow varies diurnally, and wastewater constituents will vary during the day and by building use type, which should affect biofilm stability due to variation in shear stress and solids content. Third, changing wastewater constituents, including bacterial concentrations, will affect biofilms. Donlan [37] describes the different characteristics that affect attachment of bacteria to form biofilms: substratum, conditioning, hydrodynamics, water quality, and cell properties. The change in the bacterial communities after switching to effluent water in this study highlights the importance of considering wastewater chemistry and microbiology. Changes in shear stress and water chemistry significantly impact biofilms [29, 35]. Longer-term studies of biofilm development and changes in established biofilms under varying flow rates, chemical conditions, and CROs are needed to account for these confounding factors.

## 5.0 CONCLUSIONS

The purpose of this study was to characterize interactions between biofilms and flowing wastewater by examining the transport and persistence of CROs and resistance genes. We showed that sewer biofilms form rapidly, incorporate resistant bacteria within minutes, retain them for extended periods, and subsequently release them back into the wastewater and to downstream biofilms. The following key conclusions can be drawn:

- Biofilm formation with stable bacteria and gene concentrations occurs within 2 to 10 days on previously sterile PVC, though the metagenomic profile of the communities continues to transform over two to three weeks.
- Carbapenem-resistant, planktonic bacteria in wastewater attach to established, complex sewer biofilms within hours, reaching peak concentrations within 6 days. Similarly, resistant and total bacteria, and 16S rRNA and CP-CRE gene concentrations in complex, established biofilms without target carbapenemase genes increased within hours of exposure to grit chamber wastewater. Further, they did not fluctuate more than 1 log unit/cm^2^ after 10 days of exposure to grit chamber water. Metagenomic analyses showed ongoing changes in the microbial community from 15 to 21 days.
- Bacteria and genes detached from high concentration “source” biofilms on PVC coupons immediately after water flow starts, increasing in the water to 5 log MPN/mL and 9 log GC 16S rRNA/mL, respectively, before decreasing over time.

Consideration of sewer biofilm dynamics is essential for accurately interpreting bacterial and gene concentrations in wastewater. Further investigation into the effects of hydraulic shear, total suspended solids in wastewater, bacterial species and concentrations, and sewer materials is needed to better understand biofilm dynamics. Together, these results demonstrate that sewer biofilms fundamentally transform wastewater signals through rapid uptake, prolonged retention, and delayed release, requiring a paradigm shift in how microbial indicators are interpreted in wastewater-based epidemiology and improved models that incorporate the effects of biofilms. For clinical and infection-prevention practice, these findings suggest that wastewater and sewer biofilm monitoring should be interpreted alongside patient-level data and plumbing system context, rather than as real-time proxies for active infection burden.

## DECLARATION OF GENERATIVE AI USE

During the preparation of this manuscript, the authors used ChatGPT 5.2 to improve grammar and clarity. After using this tool, the authors carefully reviewed and edited the text and take full responsibility for the final content.

## ACKNOWLEDGMENTS

This work was supported by CDC contract #200-2021-12774, Safety and Healthcare Epidemiology Prevention Research Development (SHEPheRD) 2022 Domain 1-A004: Wastewater surveillance approaches for antimicrobial resistant genes and organisms in healthcare settings within the Western U.S. Region, Jennifer Weidhaas, Ph.D. (Civil and Environmental Engineering), Principal Investigator.

The authors acknowledge the valuable assistance of Aiden Bethune (University of Utah) and Angela Coulliette-Salmond, Amanda K Lyons, Florence Whitehill, Joe Sexton, Jorge Chavez, and Margaret Williams (Centers for Disease Control and Prevention). The authors gratefully acknowledge the water treatment utility that allowed us to deploy in-line bioreactors at their facility in support of this study.

## AUTHOR CONTRIBUTIONS

**Ean Warren:** Data curation, Formal analysis, Investigation, Methodology, Visualization, Writing-Original draft and Editing. **William Brazelton:** Conceptualization, Data curation, Formal analysis, Writing-Review and Editing. **Sydney Fusco:** Data curation, Investigation, Writing-Original draft. **James VanDerslice:** Writing-Review and Editing. **L. Scott Benson:** Writing-Review and Editing. **Windy Tanner:** Writing-Review and Editing. **Jennifer Weidhaas:** Conceptualization, Formal analysis, Funding acquisition, Methodology, Project administration, Supervision, Visualization, Writing-Review and Editing.

## SUPPLEMENTAL INFORMATION

### Supplemental materials and methods Study materials

Hexadecyltrimethylammonium bromide (≥99 percent, CTAB), tris(hydroxymethyl)aminomethane buffer solution (pH 7.4, Tris), phenol:chloroform:isoamyl alcohol (25:24:1, saturated with 10 mM Tris, pH 8.0, 1 mM EDTA, PCI), chloroform–isoamyl alcohol mixture (24:1, ≥99.5 percent, BioUltra grade), meropenem were obtained from Sigma-Aldrich (St. Louis, Missouri, USA). Phosphate-buffered saline (10x Solution) was purchased from Fisher BioReagents (Fisher Scientific International, Fair Lawn, New Jersey, USA) and diluted with molecular-grade water to 1x (PBS). Polyethylene glycol 6000 (Ultrapure, PEG) was purchased from Thermo Scientific Chemicals (Fair Lawn, New Jersey, USA). Sodium chloride (Molecular Biology Reagent) was supplied by MP Biomedicals (Solon, Ohio, USA), and ethanol (200 proof) was obtained from Decon Laboratories, Inc. (King of Prussia, Pennsylvania, USA). All reagent dilutions and solutions were prepared using DNA-free water (Fisher BioReagents, Fisher Scientific International).

Bacterial cultures were obtained from CDC and Food and Drug Administration (FDA) Antibiotic Resistance Isolate Bank. Four isolates containing carbapenemase genes (*bla*_KPC_, *bla*_VIM_, *bla*_NDM_, and *bla*_IMP_) were cultivated for use in this study (**supplemental Table S2**).

Sewer biofilms were simulated using Bio-inLine Biofilm Reactors (Biosurface Technologies, Bozeman, Montana, hereafter reactors) containing 12 nylon, 2.1-cm diameter screw-in plugs with PVC coupons (1.3-cm diameter, Biosurface Technologies). The reactors are 61 cm long with a 1.6 cm^2^ square cross section and 98.3 ml working volume. Pumps include Masterflex L/S Compact Variable-Speed Drive, Masterflex L/S Digital Dispensing System, and Fisherbrand GP1000 General-Purpose Peristaltic Pumps, all calibrated to dispense 10 L/hour (170 mL/min). A flow of 10 L/hour was selected to keep the shear stress approximately the same between the square biofilm reactors and a 20.3 cm (8 inch) diameter sewer pipe sloped at 1% and flowing with 0.6 cm (0.25 inch) of wastewater at 0.31 m/s (1 ft/s) (see **below** for shear stress calculations).

### Estimation of shear stress in sewer from a single building with 8-inch PVC sewer pipe

Shear stress in the sewer was modeled using equation ES1 below:

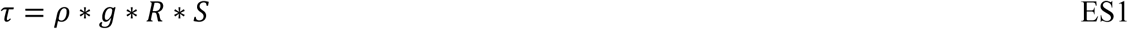

Where *τ* is shear stress in (kg/m/s^2^ or Pa); *ρ* is fluid density (1.0176 kg/m^3^); *g* is the acceleration due to gravity (9.81 m/s²); *R* is the hydraulic radius, in meters; and *S* is the slope of the channel, dimensionless. Here, the slope was assumed to be 0.5% to 1%, which falls within the typical range of 0.5% to 10%, where 1% is the optimum design slope.

The hydraulic radius *R* is estimated according to equation ES2 below:

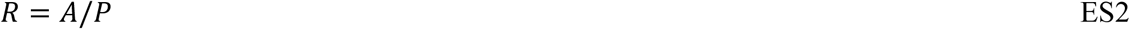

Where *A* is the cross-sectional area of flow, in m^2^; and *P* is the wetted perimeter, in m.

The cross-sectional area A of a partially full flowing pipe is estimated according to equation ES3 below:

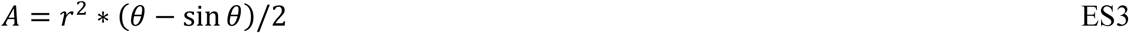

Where *r* is the radius of the pipe, in m, here assumed to be 0.203 m (8 inches); and θ is the angle from the center of the pipe to the water surface at each edge of the pipe, in degrees.

The *θ* is estimated according to equation ES4 below:

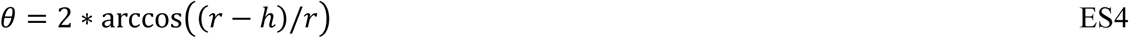

Where *h* is the depth of the water in the sewer pipe, in m, here assumed to be 0.06 m (0.25 inch). Observations at several sewer manholes collected wastewater from a single building with 100 to 300 person occupancies in 8-inch sewer pipes were observed to have typical water depths from 0.06 to 0.12 m typically (e.g., 0.25 inches to 0.5 inches water depth).

Using these assumptions, the shear stress estimated with ES1 is 0.21, 0.42, and 4.16 mPa for sewer pipe slopes of 0.5%, 1%, and 10%, respectively. This range of shear stresses can then be used to estimate the velocity required in a rectangular channel to have a similar shear stress.

### Estimation of shear stress in rectangular channel

The Darcy-Weisbach equation for shear stress under pumping conditions in a full-flowing rectangular channel is described in equation ES5 as follows:

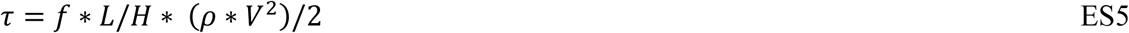

Where *f* is the friction factor that depends on the Reynolds number *Re* and is calculated as shown in ES6; *L* is the length of the channel, in m; *H* is the hydraulic diameter, in m; and *V* is the water velocity in the channel, in m/s. In square channels used in the reactor herein with a cross-sectional area of 1.61 cm^2^, the hydraulic diameter, *H*, is 0.052 m. The length of the reactor, *L*, is 61 cm.

The following equation ES6, the Moody’s approximation, can be used to estimate *f* for pipes with a Reynolds number (*Re*) in the range of 4*10^3^ to 4*10^8^.

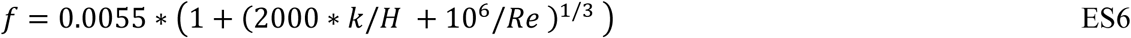

Where *k* is the surface roughness, in m. The *Re* range was 1000 to 10,000 in the laminar to turbulent flow regime. The relative roughness is the ratio of the absolute roughness to the hydraulic diameter. As the absolute roughness of new and aged PVC ranges from 0.0015*10^−3^ to 0.07*10^−3^ m, and for rusted steel ranges from 0.15*10^−3^ to 4*10^−3^ m. Here we used a range of relative roughness for biofilms in sewers from 0.07*10^−3^ to 4*10^−3^. The estimated coefficient of friction then ranges from 0.031 to 0.07. Solving ES5 for velocities to match the range of shear stresses in the sewer pipe shows that velocities fall in the range of 0.022 to 0.047 m/s. These velocities use the 0.06 m water flow depth, 0.5% to 1% slopes, and *Re* of 1,000 to 10,000. For this square channel, the volumetric discharge is 13 to 27 L/hour. If we assumed a 10% slope, the volumetric discharge required to match the sewer shear stress was 86 L/hour. Therefore, using 10 L/hour in our studies is either approximately equal to the lowest estimated shear stress in the sewer or is 11% of the maximum likely shear stress in the sewer.

### Meropenem in grit basin and facility effluent water

To measure meropenem concentrations, one-liter samples of grit basin and facility effluent water were collected. Meropenem was extracted from grit basin water by solid-phase extraction (SPE) method. One-hundred fifty mL of sample in three equal aliquots was centrifuged at 15,500 *x g* for 20 minutes. The supernatant was filtered through a 934-AH glass fiber filter (Whatman, Maidstone, United Kingdom). The filtrate was filtered again through a 0.45 µm mixed ester cellulose membrane filter (Fisher Scientific International, Fair Lawn, New Jersey, USA). The SPE method followed the recommendations of Sharma 1. In brief, Oasis HLB 500 mg LP Extraction cartridges (Waters Corporation, Milford, Massachusetts, USA) were conditioned with 5 mL of 100% HPLC-grade methanol (Fisher Scientific International), followed by 5 mL of HPLC-grade water (Fisher Scientific International). One hundred mL of filtrate was loaded onto the Oasis cartridge. The cartridge was rinsed with 1 mL of water, followed by 1 mL of 5% methanol. The cartridge was dried under vacuum for 5 minutes, then rinsed with 1 mL 70% methanol. Meropenem was eluted with the subsequent 70% methanol. Facility effluent water sample was centrifuged (15,500 *x g* for 20 minutes) then triple-filtered with a (1) 934-AH glass fiber filter, (2) 0.45 µm mixed ester cellulose membrane filter, and (3) 0.22 µm glass fiber filter (Tisch Scientific, Cincinnati, Ohio, USA). Duplicates of both samples were spiked with 10 mg/L meropenem.

Samples were analyzed using an Agilent (Santa Clara, California, USA) 6470 Triple Quadrupole liquid chromatography mass spectrometer with a guard column (SecurityGuard ULTRA Cartridges for C18 UHPLC column, Phenomenex, Torrance, California, USA) and an Acquity UPLC HSS T3 1.8 µm, 2.1 × 100 mm column (Waters Corporation). The eluents were composed of 0.1% formic acid (LiChropur, Millipore, Burlington, Massachusetts, USA) in LCMS-grade water (Fisher Scientific) and acetonitrile at a flow rate of 0.3 mL/minute with proportions shown in **supplemental Table S1**. The mass spectrometer settings were as follows: electrospray ionization in positive mode, spray voltage 3800 V, gas temperature 300 °C, gas flow 8 L/min, nebulizer 35 psi, sheath gas temperature 300 °C, sheath gas flow 11 L/min, precursor ion 384.2, production ion 68.1, fragmentor 135, collision energy 35, and cell accelerator voltage 4 V.

### Biofilm total and volatile solids

To measure total and volatile solids, coupons with biofilms were sonicated on high in 10 mL deionized water for 10 minutes at 37 °C [2]. The water was transferred to pre-weighed aluminum pans and dried at 105 °C for two hours. The dried pans were weighed to measure total solids. The solids were heated at 500 °C for 15 minutes to measure volatile solids. Grit found on coupons was similarly treated, except without the sonication step.

### Quantification of total and resistant bacteria

Biofilm bacteria on PVC coupons were measured by scraping the diameter twice with sterile pipette tips. The tips were rinsed five times in 1 mL of sterile PBS to suspend the bacteria for later dilution. Total bacterial counts were determined using LB agar plates or three-tube most probable number assay (MPNs) with Tryptic Soy Broth (TSB) or LB media. CROs were determined using three-tube MPNs with 8 mg/L meropenem or imipenem or on CHROMagar KPC. MPNs were performed in 96-well plates, and bacterial presence in the wells was measured on a Biotek Synergy HTX (Agilent, Santa Clara, California) microplate reader at OD600.

### Nucleic acid extraction

Nucleic acid extraction of coupons, influent grit basin water, and effluent reactor water was performed by following a previously published method [3] with the following modifications. Coupon nucleic acids were extracted by placing the coupons in 5 mL PowerWater DNA Bead Tubes (QIAGEN, Hilden, Germany) containing 1.25 mL PCI and 1.25 mL CTAB. The tubes were mixed by vortex for 10 minutes at setting 8 and then centrifuged at 3,000 *x g*. The supernatants were transferred to 2 mL centrifuge tubes for a chloroform:isoamyl alcohol cleaning step. The extracts were incubated with 30 percent PEG and sodium chloride for two hours then centrifuged twice for 15 minutes at 17,000 *x g* with a 100 µL 70 percent ice-cold ethanol wash between spins. The resultant pellets were dissolved in 50 µL 10 mM Tris for subsequent quantitation by quantitative polymerase chain reaction (qPCR).

Effluent water samples from the reactors were filtered through sterile 0.45 µm mixed cellulose ester membranes (Fisher Scientific International, Fair Lawn, New Jersey, USA). The filters were cut in half with bleach-sterilized scissors and processed similarly to the PVC coupons, with the bacteria-facing side of the filter facing the beads in 5 mL PowerWater DNA Bead Tubes. The rest of the method followed the extraction described above.

Nucleic acids from grit basin water used as influent to the reactors were extracted by aliquoting 40 mL into centrifuge tubes [see 4]. One percent PEG and 0.9 percent sodium chloride were added, and the tube was shaken at 4 °C for two hours. The tube was centrifuged for 10 minutes at 31,400 *x g*. The supernatant was decanted, and the pellet was suspended in sterile 1.5 mL PBS. Five hundred µL was added to Lysing Matrix E bead tubes (MP Biomedicals, Eschwege, Germany) with 500 µL PCI and 500 µL CTAB and mixed by vortex for 10 minutes at setting 8. The rest of the method followed the extraction described above.

### Metagenomics

Metagenomic sequencing of extracted DNA was conducted at Utah Public Health Laboratory with an Illumina NextSeq platform (P2 flow cell, 300-cycle kit for 2 x 150 bp paired reads). Metagenome sequences were obtained from 55 biofilm samples from Experiments 1, 2, and 3 (**Table 1**). Six of these samples yielded <50,000 reads per sample and were not included in downstream analyses. An additional 15 samples yielded less than 50 million sequences per sample; these were included in downstream analyses but removed when plotting alpha diversity results. The remaining 34 samples yielded 50 - 120 million reads per sample. Adapter sequences and PhiX were removed from all reads with BBDuk (part of the BBTools suite, v39.28 [5] and quality trimming was performed with fastp v1.0.1 [6] using a sliding window of eight bases and a quality threshold of 28. Ninety-five to ninety-eight percent of reads and bases were retained after quality filtering and trimming. Reads from each sample were assembled with Megahit v1.2.9 [7], and genes were predicted with Prodigal v2.6.3 [8] in meta mode.

Quality-filtered, unassembled reads were assigned taxonomy with singlem v0.19.0 [9] and the GTDB taxonomy database r226 [10]. singlem uses a marker gene approach to identify metagenomic operational taxonomic units (OTUs) that can be used for standard diversity analyses. The number of metagenomic OTUs in each biofilm sample was correlated with sequence depth when including all 49 samples that were successfully sequenced. After removing 15 samples with <50 million reads per sample, no correlations between sequence depth and alpha diversity metrics remained among the remaining 34 biofilm metagenomes. Therefore, the set of 34 metagenomes with high sequence coverage was used to report alpha and beta diversity results (i.e., numbers of OTUs and genes, and multivariate analyses of OTU and AMR gene compositions).

Antimicrobial resistance genes (AMR genes) were identified among predicted proteins in metagenomic assemblies from the set of 49 metagenomes with AMRFinderPlus v4.0.23 [11]. When reporting results, AMRFinder hits labeled as HMM or PARTIALP were excluded. The metagenomic sequence coverage of each AMR gene is reported as the coverage of the assembled contig that encodes it, calculated with CoverM v0.7.0 [12]. Coverages of AMR genes are reported at the “Element symbol” level, as defined by AMRFinderPlus. Coverages are reported in units of transcripts (or fragments) per million (TPM). TPM is a proportional unit (multiplied by one million for convenience) that is normalized to the length of each contig as well as to the total library size.

Taxonomic classification results and AMR gene results were plotted as non-metric multidimensional scaling (NMDS) plots using Bray-Curtis dissimilarity scores and the metaMDS command within the vegan package v2.7-1 in R [13].

### Mass balance for Experiment 5

The compartments assessed were (1) the most probable number (MPN) or CP genes in the established biofilm prior to exposure to the isolates as shown in Equation 1, (2) the MPN or carbapenemase genes on the source coupons dipped in the isolate culture and rinsed with PBS as shown in Equation 2, (3) the MPN or carbapenemase genes on the source coupon after the 1 hour of water flow as shown in Equation 3 and (4) the cumulative effluent MPN or carbapenemase genes over the 1 hour of water flow shown in Equation 4.

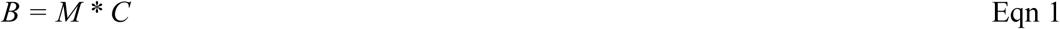

Where *B* is the background mass of bacteria or genes, *M* is the MPN/cm^2^ or carbapenemase genes/cm^2^ on the PVC coupon after removal from the sewer, and *C* is the PVC coupon surface area in cm^2^.

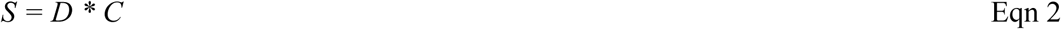

Where *S* is the starting mass of bacteria or genes, *D* is the MPN/cm^2^ or carbapenemase genes/cm^2^ on the position 1 PVC coupon after dipping the coupon in the isolate culture for 2 minutes, followed by rinsing.

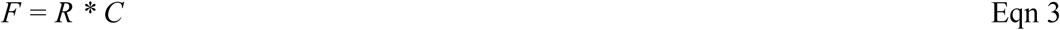

Where *F* is the final mass of bacteria or genes, *R* is the MPN/cm^2^ or carbapenemase genes/cm^2^ on the position 1 PVC coupon after one hour of water flow through the reactor.

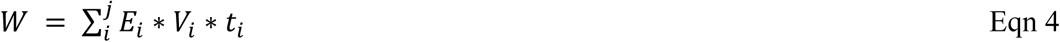

Where *W* is the mass of bacteria or genes leaving the reactor in the water; *i* is the step increment until the last increment, *j*; *E_i_* is the reactor effluent MPN/mL or carbapenemase genes/mL in each increment *i*; *V_i_* is the ml of water collected in each time increment *i*; and *t_i_* is the minutes for each increment i. The total mass of MPN or carbapenemase genes is then the summation as shown in Eqn 5.

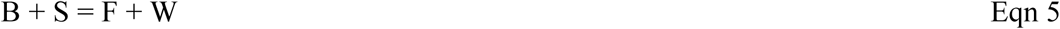

### Supplemental results

#### Grit basin and facility effluent water meropenem concentrations

Standard curves were linear (r^2^ = 0.999) over the range of 0.01 to 0.20 µg/mL for meropenem. Recovery of the meropenem in grit basin water using the SPE method was 24% and 29% using the filtration method for the effluent water. Meropenem was not detected in either the grit basin or the effluent water at 0.01 µg/mL, indicating that the sample concentrations, after accounting for extraction recoveries, were less than 0.08 µg/mL. This concentration is at least 10 times less than the minimum inhibitory concentrations (MICs) of meropenem in four strains of planktonic *Acinetobacter baumannii* [14]. *A. baumannii* in biofilms were more resistant with minimum biofilm eradication concentrations of 256 to 4096 µg/mL. Susceptible *Klebsiella pneumoniae* without carbapenemase genes were resistant at 1-2 µg/mL of meropenem [15]. These results indicate there is no selective pressure to maintain meropenem resistance.

---

**Supplemental Table S1.** Eluent proportions for meropenem determination by liquid chromatography.

| Time,<br>minute | 0.1% Formic<br>acid, % | Acetonitrile,<br>% |
| --- | --- | --- |
| 0.0 | 99 | 1 |
| 1.0 | 75 | 25 |
| 5.0 | 10 | 90 |
| 5.5 | 1 | 99 |
| 6.5 | 99 | 1 |
| 7.0 | 99 | 1 |

**Supplemental Table S2.**
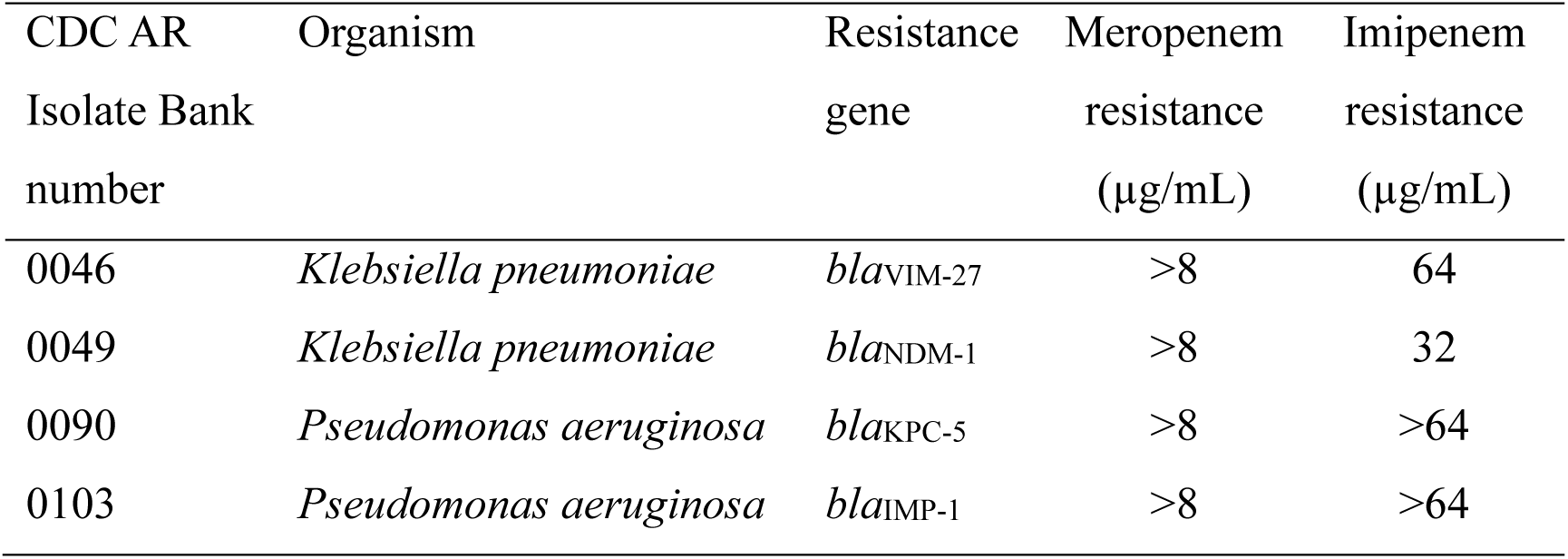
CDC and Food and Drug Administration (FDA) Antibiotic Resistance Isolate Bank numbers, species, gene, and meropenem resistance used in Experiments 4 and 5.

| CDC AR<br>Isolate Bank<br>number | Organism | Resistance<br>gene | Meropenem<br>resistance<br>(µg/mL) | Imipenem<br>resistance<br>(µg/mL) |
| --- | --- | --- | --- | --- |
| 0046 | *Klebsiella pneumoniae* | *bla*<sub>VIM-27</sub> | >8 | 64 |
| 0049 | *Klebsiella pneumoniae* | *bla*<sub>NDM-1</sub> | >8 | 32 |
| 0090 | *Pseudomonas aeruginosa* | *bla*<sub>KPC-5</sub> | >8 | >64 |
| 0103 | *Pseudomonas aeruginosa* | *bla*<sub>IMP-1</sub> | >8 | >64 |

**Supplemental Table S3.**
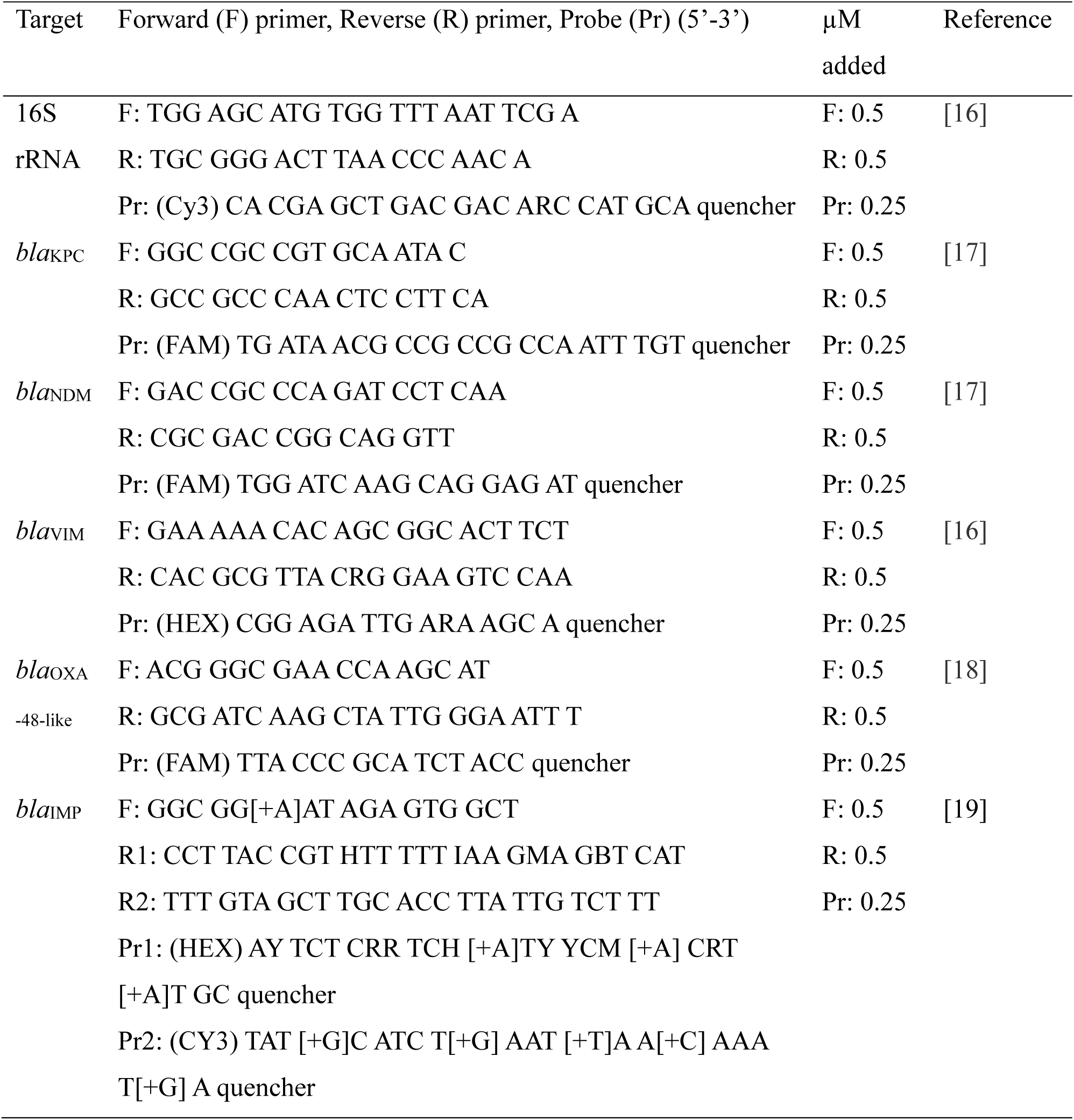
Quantitative polymerase chain reaction (qPCR) primers and probes.

**Supplemental Table S4.**
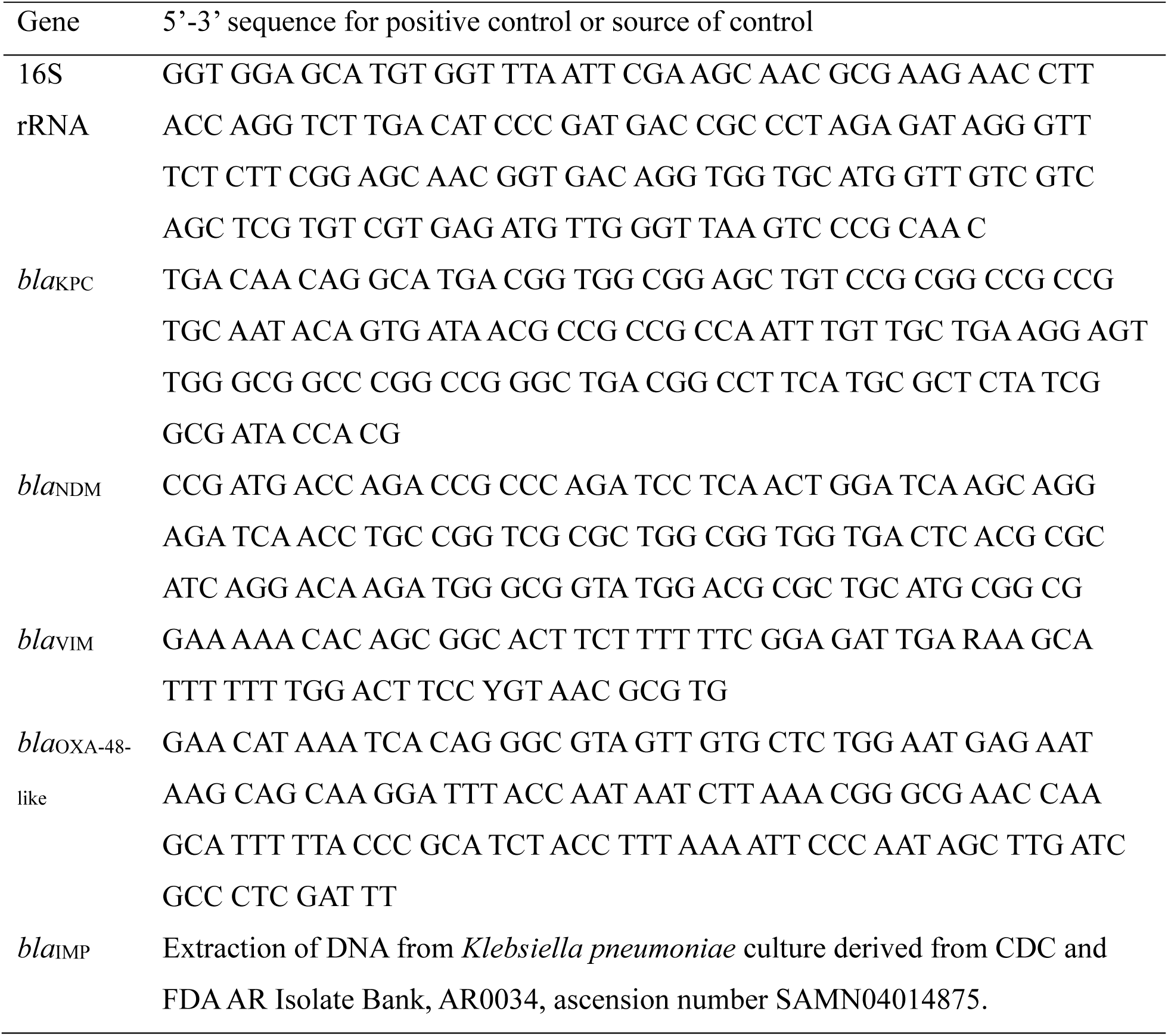
Positive controls used for quantitative polymerase chain reaction (qPCR) [4].

**Supplemental Table S5.** Quantitative polymerase chain reaction (qPCR) reaction efficiencies (%) for single-plex and duplex gene assays using a QuantStudio 3 Real-Time PCR machine (Thermo Fisher Scientific, Waltham, Massachusetts) [4]

| Single-plex (fluor) | Efficiency (%) | R<sup>2</sup> | n |
| --- | --- | --- | --- |
| Zymo (HEX) | 96.0 | 1.000 | 7 |
| KPC (FAM) | 94.0 | 1.000 | 7 |
| 16S (Cy3) | 99.1 | 0.999 | 7 |
| NDM (VIC) | 115.6 | 0.995 | 5 |
| NDM (FAM) | 98.7 | 0.997 | 7 |
| IMP (CY3) | 93.1 | 0.997 | 6 |
| VIM (HEX) | 90.5 | 0.996 | 8 |
| OXA (FAM) | 100.1 | 0.999 | 7 |
| Duplex (fluor) | Efficiency (%) | R<sup>2</sup> | n |
| Zymo (HEX) with *C. auris* | 88.9 | 0.999 | 7 |
| KPC (FAM) with 16S | 92.6 | 0.999 | 7 |
| 16S (CY3) with KPC | 119.0 | 0.991 | 7 |
| NDM (VIC) with OXA | 97.3 | 0.994 | 7 |
| NDM (FAM) with IMP | 110.5 | 0.995 | 6 |
| IMP (CY3) with NDM | 96.3 | 0.994 | 6 |
| VIM (HEX) with OXA | 96.7 | 0.995 | 7 |
| OXA (FAM) with NDM | 97.8 | 0.999 | 7 |
| OXA (FAM) with VIM | 97.8 | 1.000 | 7 |

**Supplemental Table S6.** Biofilm reactor influent parameters with standard deviations for Experiments 1, 2, and 3. “n” represents the number of samples analyzed.

| Parameter | Grit basin wastewater<br>1 Jul 2025 – 22 Jul 2025 | Effluent water<br>27 Jul 2025 – 5 Aug 2025 |
| --- | --- | --- |
| Temperature, °C | 21.05 ± 0.67 (n=22) | 22.87 ± 0.68 (n=10) |
| pH | 7.51 ± 0.13 (n=22) | 7.19 ± 0.04 (n=10) |
| Carbonaceous Biochemical |  |  |
| Oxygen Demand, mg/L | 195.64 ± 33.52 (n=14) | 5.80 ± 1.49 (n=6) |
| Total suspended solids, mg/L | 252.38 ± 72.22 (n=16) | 27.40 ± 2.40 (n=9) |
| Total Kjeldahl Nitrogen, mg/L | 47.67 ± 2.08 (n=3) | 0.83 ± 1.03 (n=2) |
| Total phosphorus, mg/L | 5.51 ± 0.09 (n=3) | 3.65 ± 0.44 (n=2) |
| Conductivity, µS/cm | 1378 ± 229 (n=22) | NA |

## Supplemental figures

**Supplemental Figure S1.**
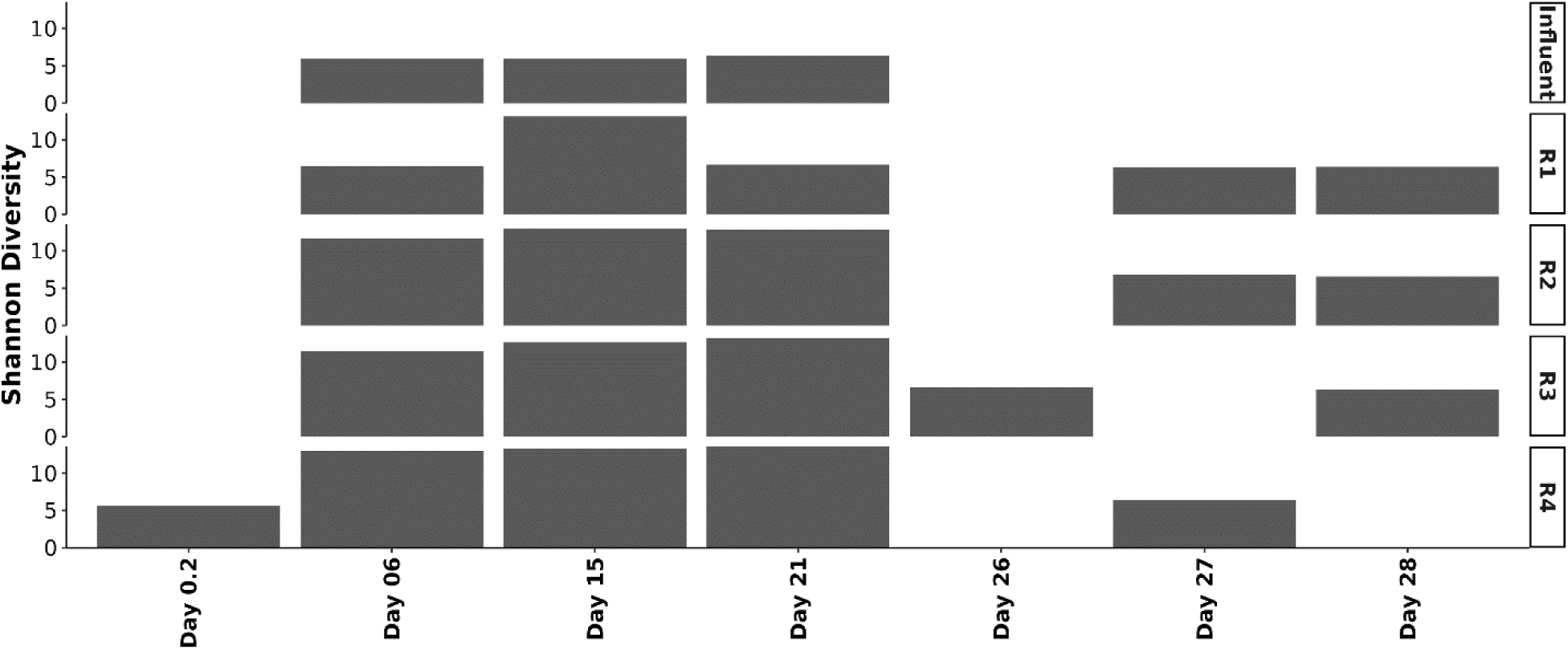
Metagenomic OTU Shannon diversities. R1 through R3 represent triplicate reactors in Experiments 2 and 3. R4 is the single reactor in Experiment 1.

**Supplemental Figure S2.**
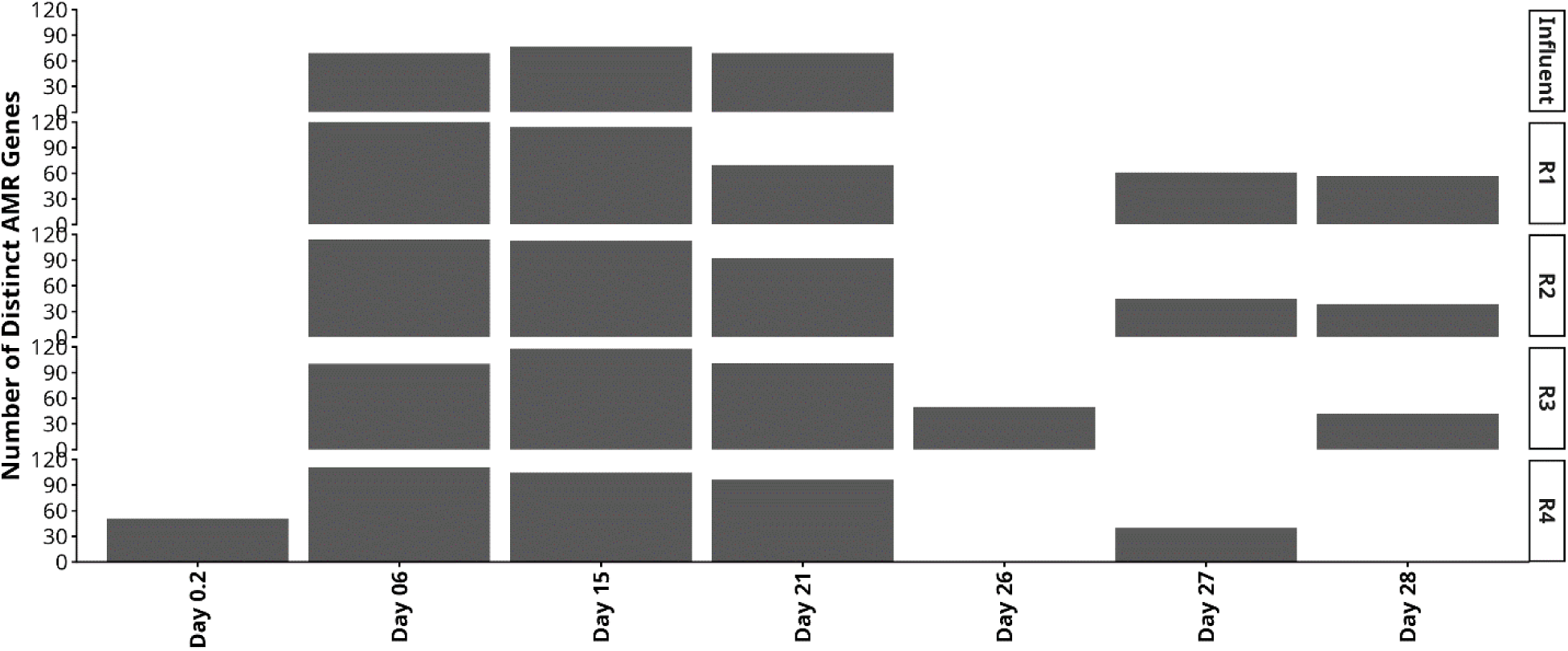
Number of distinct Antimicrobial Resistant (AMR) genes in the biofilm metagenomes, where a distinct gene is defined as an “Element symbol” identified by AMRFinder.

